# Epithelial zonation along the mouse and human small intestine defines five discrete metabolic domains

**DOI:** 10.1101/2023.09.20.558726

**Authors:** Rachel K. Zwick, Petr Kasparek, Brisa Palikuqi, Sara Viragova, Laura Weichselbaum, Christopher S. McGinnis, Kara L. McKinley, Asoka Rathnayake, Dedeepya Vaka, Vinh Nguyen, Coralie Trentesaux, Efren Reyes, Alexander R. Gupta, Zev J. Gartner, Richard M. Locksley, James M. Gardner, Shalev Itzkovitz, Dario Boffelli, Ophir D. Klein

## Abstract

A key aspect of nutrient absorption is the exquisite division of labor across the length of the small intestine, with individual classes of micronutrients taken up at different positions. For millennia, the small intestine was thought to comprise three segments with indefinite borders: the duodenum, jejunum, and ileum. By examining fine-scale longitudinal segmentation of the mouse and human small intestines, we identified transcriptional signatures and upstream regulatory factors that define five domains of nutrient absorption, distinct from the three traditional sections. Spatially restricted expression programs were most prominent in nutrient-absorbing enterocytes but initially arose in intestinal stem cells residing in three regional populations. While a core signature was maintained across mice and humans with different diets and environments, domain properties were influenced by dietary changes. We established the functions of *Ppar-ẟ* and *Cdx1* in patterning lipid metabolism in distal domains and generated a predictive model of additional transcription factors that direct domain identity. Molecular domain identity can be detected with machine learning, representing the first systematic method to computationally identify specific intestinal regions in mice. These findings provide a foundational framework for the identity and control of longitudinal zonation of absorption along the proximal:distal small intestinal axis.

## Introduction

In the small intestine, regional specialization optimizes digestion by enabling distinct micronutrients to be sequentially absorbed at different anatomical positions. Traditionally, the small intestine has been separated into three loosely defined regions: the duodenum, jejunum, and ileum. These segment designations, which date back to observations made by the ancient Greeks, are thought to correlate with various absorptive processes, but their anatomical boundaries are vague^1^. In addition to differences in tissue structure and cellular composition along the length of the intestinal epithelium to support specialized functions, many genes show variable spatial expression patterns, as recently illustrated by single-cell RNA sequencing (scRNAseq) comparisons of epithelial cells from the classical regions of the mouse and human small intestine and colon^2-7^. However, apart from the human duodenojejunal flexure, which is suspended by the ligament of Treitz, a lack of discrete landmarks to anchor these regional definitions precludes examination of the precise organization and properties of local niches within the small intestine. The extent to which the three classical parts of the small intestine explain the complexity of regional patterns in the tissue, and how these patterns respond to environmental changes such as nutrient fluctuations, pathogen exposures, and disease, is not clear.

By contrast with the mammalian small intestine, the *Drosophila* midgut divides into 10- 14 distinct compartments, of which a subset have been shown to contain intestinal stem cells (ISCs) with innate regional properties^8-11^. These findings raise the possibility that mammals may exhibit more finely grained intraintestinal spatial differences than have been appreciated and that adult intestinal stem cells (ISCs) may program functional environments within the tissue. In line with the latter possibility, regional expression of numerous genes, including those associated with absorption, is maintained in mouse and human intestinal organoid cultures *ex vivo*^12-14^. However, the molecular programs encoded in ISCs that specify the expression of regionalized functional genes in their differentiated progeny are not known.

Here, we report the transcriptional programs, associated metabolic functions, and locations of five previously undefined epithelial regions within the mouse and human small intestine. We track the refinement of regional patterns across the absorptive lineage from ISCs to specialized enterocytes and establish a cellular and molecular model explaining how they are maintained by epithelial-intrinsic mechanisms throughout adulthood.

## Results

### Five groups of enterocytes occupy distinct zones along the proximal to distal length of the mouse and human small intestine

To study the mechanisms that maintain intestinal regionality, we took an unbiased approach to define the organization of the intestine on a molecular level, asking: how many functional domains, defined by distinct cellular states, are present in the mammalian small intestine? While previous studies of regional identity assumed the presence of three major regions – the duodenum, jejunum, and ileum – and sampled the intestine to best approximate their positions^2-7^, we set out to examine the small intestine without preconceptions. Our approach leveraged MULTI-seq scRNAseq multiplexing^15^ to barcode cells collected from 30 equally sized segments spanning the entire length of the small intestines of both mouse and human (Fig. 1a). We used tissue from two Lgr5-GFP mice in which stem and progenitor cells – ISCs and their immediate transit amplifying (TA) cell progeny – express GFP, and from two human donors. We sequenced total epithelial cells (CD45^-^, pan-epithelial EpCAM^+^) and an equal number of progenitor cells (crypt marker CD44^+^ in mouse and human cells, Lgr5-GFP^+^ in mouse cells). We recovered a total of 19,847 mouse cells and 36,588 human cells (Fig. 1a, Extended Data Fig. 1-5, and Methods), including all progenitor and specialized intestinal epithelial cell types (Fig. 1, b and c, Extended Data Fig. 6), aside from CD45+ tuft cells^2^.

**Fig. 1.**
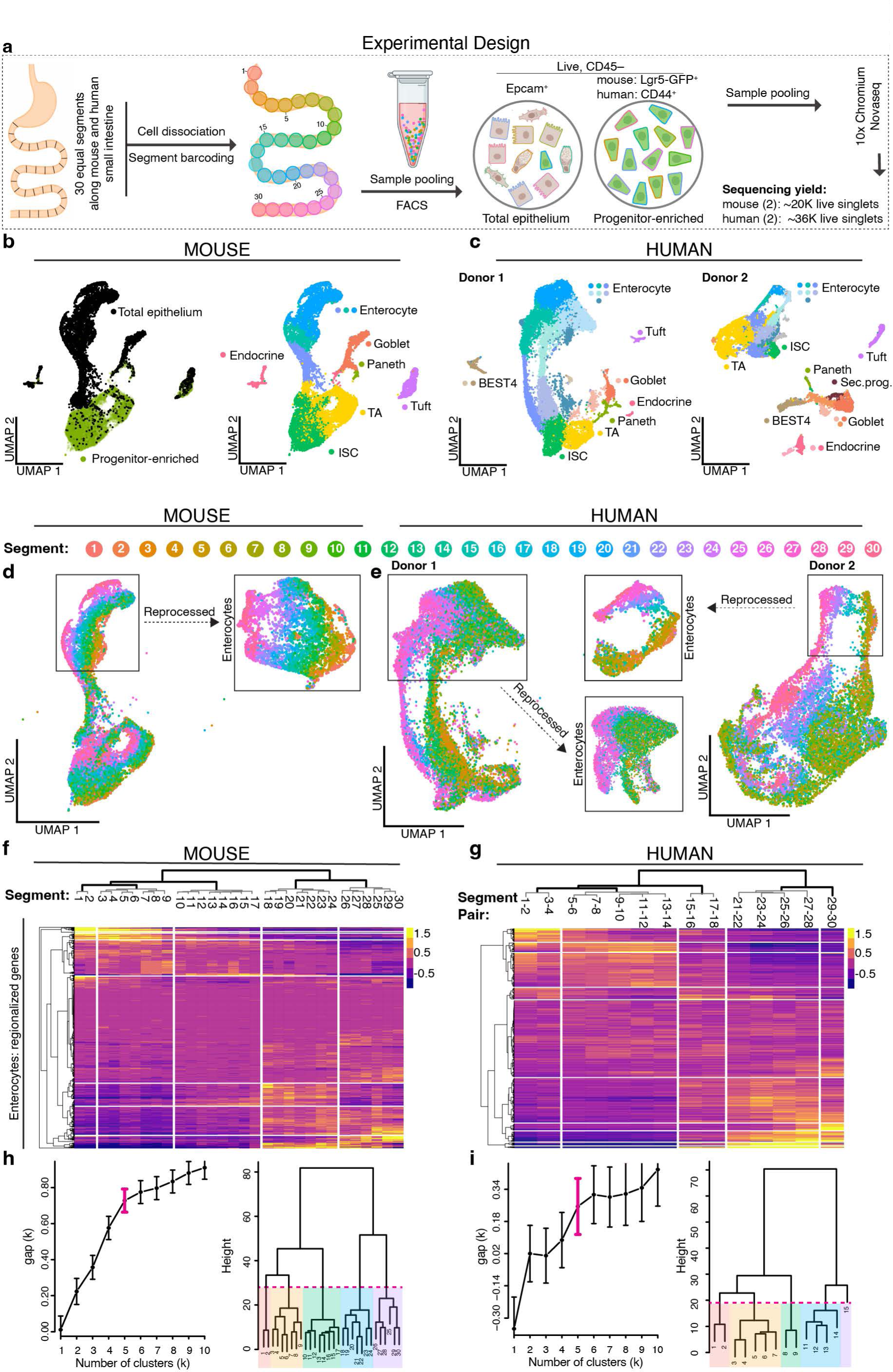
Five groups of enterocytes occupy distinct zones along the proximal to distal length of the mouse and human small intestine. **a** Schematic of the strategy for scRNAseq of epithelial cells from 30 equal segments of the mouse (n = 2) and human (n = 2) small intestine. Cells from each segment were dissociated, tagged with segment-specific barcodes, pooled, sorted into total epithelial and progenitor-enriched samples, and sequenced. Cell number yields following data QC are shown. **b,c** UMAP of sequenced mouse and human cells following QC, annotated with total epithelial or progenitor-enriched sample identification (b, left) or predicted cell type. M-cells not displayed, c.f. Extended Data Fig. 5–7. **d,e** UMAP of absorptive lineage cells colored by segment number along the proximal to distal axis. Insets display reprocessed enterocyte subsets. Human donor 2 is used for subsequent main figure panels unless otherwise noted. **f,g** Average expression of the top 150 upregulated genes in mouse and human enterocytes in each segment, with segment order and hierarchical clustering based on expression distance between segments. Vertical white lines show the five domains that divide the small intestine, based on: **h,i** *left:* gap statistics for hierarchical clusters of enterocytes in regional gene expression distance. *Right:* Cuts of dendrograms with optimal cluster numbers (magenta brackets, left), with the branches and segment numbers of five resulting regional enterocyte groups shaded. UMAP: Uniform Manifold Approximation and Projection.

Visualization of the 30 segments in gene expression space for mouse and human scRNAseq data revealed pronounced shifts in cell state along the proximal:distal axis (Fig. 1, d,e). While regionally variable genes were evident in all epithelial cell types, including secretory cells (Extended Data Fig. 7a, Supplementary Table 1), such shifts were most stark in enterocytes, with > 80% of genes expressed by these cells in mouse and human being significantly zonated along the longitudinal axis (q < 0.05 using Kruskal-Wallis test on genes with mean sum-normalized expression above 5 X 10^-6^). In the mouse intestine, vertical zonation from the crypt/villus base boundary to the tip of the villus, previously studied only in the jejunum^16^, was maintained across the proximal:distal axis (Extended Data Fig. 7b-e). These data demonstrate the impact of cell position along multiple axes on enterocyte gene expression.

We next asked whether transcriptional progression along the proximal:distal axis of the small intestine is continuous, or if and where discontinuous transitions in gene expression divide the duodenum, jejunum, and ileum and/or an alternative set of regions. Focusing on enterocytes, which were the most highly zonated epithelial cell type, we computed the average expression of the 150 most regionalized genes in enterocytes from each segment and performed hierarchical clustering on the resulting data (Fig. 1f,g and Extended Data 8a). Remarkably, this computational approach reconstructed the anatomical order of segments in the mouse small intestine with almost perfect accuracy (cf. segment numbers in dendrogram Fig. 1f, where all segments are in the correct numerical order apart from segments 14-16 and 25), reinforcing the primacy of regional position in defining enterocyte transcriptional states. We also observed essentially perfect ordering of human segments, which were grouped into pairs due to cell number variability by segment (cf. segment pair numbers in dendrogram Fig. 1g, ordered accurately except for the missing pair 19-20, from which insufficient cell numbers were captured in the displayed sample).

The computational approach used to order segments was also used to define their higher-level organization. Specifically, the Euclidian distance between enterocyte gene expression in individual segments measured which segments had most similar expression profiles and clustered them accordingly. The resulting hierarchical clusters (dendrograms, Fig. 1f,g, Extended Data Fig. 8a) revealed the order in which segments form groups at increasingly higher levels. We used the gap statistic to estimate the optimal number of enterocyte clusters^17^. In this method, gap values rise more steeply with an increasing number of well-separated clusters and rise less steeply, or remain stable, with additional unnecessary clusters. In both mouse and human, five was the peak gap value prior to a flattening of the gap statistics (magenta bracket, Fig. 1h,I and Extended Data Fig. 8b, left). Notably, the boundaries of five domains were stable when using fewer genes than 150, indicating that a five-domain superstructure is not dependent on the number of genes used for its identification (Extended Data Fig. 8c). Our clustering analysis revealed that mouse and human enterocytes optimally divide into 5 clusters of regional expression profiles, as displayed in the corresponding cuts of the dendrograms (Fig. 1h,I and Extended Data Fig. 8b, right).

We then evaluated zonal enterocyte clustering based on a second metric, Jensen-Shannon divergence (JSD). JSD provides a separate method to evaluate shifts in gene expression based on quantification of the distances between enterocytes in segments plotted by UMAP. Hierarchical clustering of the resulting distance matrix for each mouse individually provided nearly identical results to our clustering based on the expression of regional genes (Extended Data Fig. 8d). Collectively, these data establish the positions of five domains of the intestine that contain transcriptionally distinct enterocytes. We have designated these regions domains A–E. On a morphological level, we observed that domains A–D displayed significantly different villus lengths, suggesting that the overall surface area available for nutrient absorption might differ between domains (Extended Data Fig. 8e).

### A progression of five distinct gene signatures divides the intestinal proximal:distal axis

We next investigated the identity and regional expression patterns of genes that delineate domains A–E (Supplementary Tables 2 and 3). Given similarities in the number and position of domains in mouse and human, we asked whether the species might share domain-defining genes. While regional profiles of genes such as human domain A signature gene adenosine deaminase (*ADA)* differed between species, we observed correlation between many of the most highly regionalized genes in both species (Fig. 2a, RSpearman = 0.29, and see Methods). For example, expression of *Pdx1* and *Hoxb*, which encode homeobox proteins at the extreme ends of the intestines of both species, and of many genes required for nutrient processing along their lengths, suggests conserved regional specialization of tissue patterning and nutrient metabolism. We noted that several genes displayed stark restriction to a single domain (i.e. another homeobox gene *Meis2* in domain A, the ileal fatty acid binding protein *Fabp6* in domain E), whereas others had broader expression, but with peaks in a given domain (i.e. sucrase isomaltase *Sis* in domain C) (Fig. 2b and Extended Data Fig. 8f). To determine whether these individual trajectories were representative of aggregate gene expression patterns that define each domain, we plotted signature scores based on the mean scaled expression of the top 20 domain-defining genes for each domain across the length of the intestine (Fig. 2c,d). Domains A, D, and E had regionally confined scores, illustrating their distinct transcriptomic signatures. Domains A and B directly overlap in the proximal-most intestine of both species, the key difference between these domains being that a small set of unique domain A-specific genes decline sharply, whereas genes common to both groups gradually decline over a larger area. While domain C displayed the least zonated molecular profile of the five domains, reflected by the broad expression of defining genes outside of domain C, its expression pattern was clearly distinct from those in neighboring domains in both species. Domain E-associated transcripts emerge where domain D declines and maintain high expression at the extreme distal end of the small intestine.

**Fig 2.**
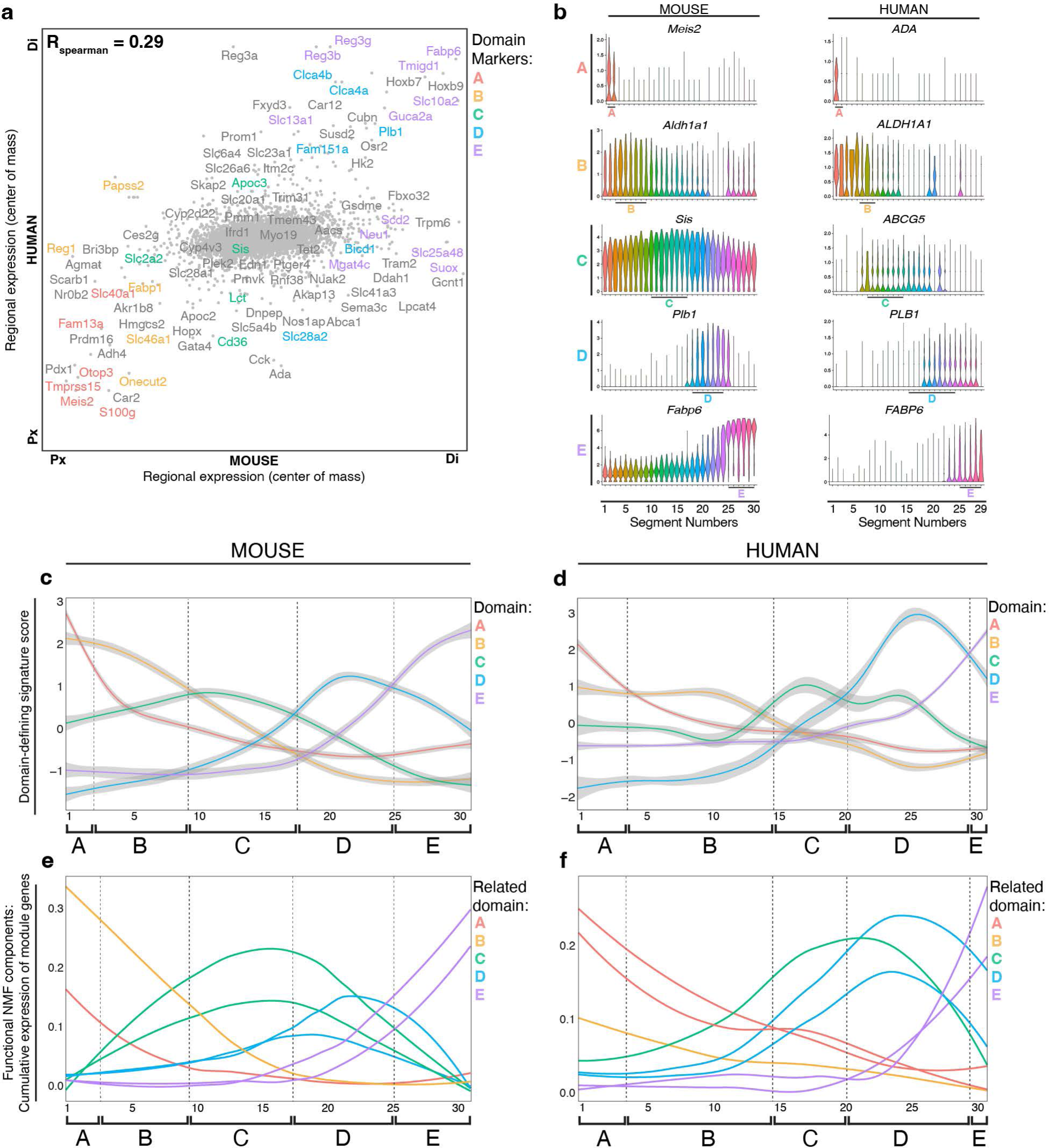
A progression of five distinct gene modules divides intestinal length. **a** Comparison of segment centers of mass for 6,191 homologous genes in mouse and human enterocytes with mean sum-normalized levels >1x10-5 in at least one point along intestinal length in both species. RSpearman = 0.29, p = 2.7 x 10-135, n = 2 mice and 2 human donors. Top segmentally variable genes in each species are shown, of which mouse domain signature genes are color-coded as indicated. Px and Di identify the proximal and distal ends of the mouse (x-axis) and human (y-axis) small intestine. **b** Expression level by segment of select marker genes of each domain in mouse and human enterocytes. Human genes were domain-enriched in both donors, representative plots from donor 1 are shown. **c, d** Domain-defining gene expression scores for mouse (**c**) and human donor 2 (**d**), which represent the mean scaled expression of the top 20 domain-defining genes, colored by domain with surrounding grey standard error bounds, across intestinal segments. Segment positions are numbered (x-axis) and positions of domain boundaries calculated in Fig. 1h,i are noted with dotted lines and brackets. **e,f** Cumulative expression of regionally variable mouse (**e**) and human (**f**) NMF gene modules across intestinal segments. Gene modules that encode physiological functions associated with nutrient metabolism are displayed. Module lines colored according to the domain A–E they most closely resemble based on regional expression trajectory and signature gene expression. Segment positions are numbered (x-axis) and positions of domain boundaries calculated in Fig. 1h,i are noted with dotted lines and brackets. NMF non-negative matrix factorization.

We then investigated the larger gene expression programs underlying the five domains reflected by their signature scores using non-negative matrix factorization (NMF). NMF detects co-expressed gene modules and, unlike the signature score approach, is agnostic to putative domain boundaries defined in our study. In both mouse and human, we detected modules that displayed variability across the small intestine (Fig. 2e, f and Supplementary Table 4). Many of these modules contained top regional signature genes (from Supplementary Tables 2 and 3), and their expression trajectories across the intestine grouped into patterns that recapitulated the signature scores. We observed two groups of components that were highly expressed at the proximal end of the intestine and declined across different breadths (as with domains A and B); components that rose and fell within the boundaries of the small intestine that organize into two groups – one that peaked around the center of the intestine and one that peaked mid-way through the distal half of the intestine (roughly within the boundaries of domains C and D); and finally components that increased concurrently with the decline of domain D-associated components and did not decline within the tissue (as with domain E). Thus, our NMF analysis reinforced the presence of five major patterns of regional gene expression by enterocytes across the intestine.

### Domain identity can be detected across samples and used for systematic classification of intestinal regions

We used multiplexed single-molecule *in situ* hybridization to validate domain assignments by probing multiple regional signature genes across coiled, full-length murine intestinal tissue (Fig. 3a and Extended Data Fig. 9-10) and human tissue collected from precise positions (Fig. 3b,c and Extended Data Fig. 11). In mice, segregated localization was observed for the domain A and D markers *Meis2 and Plb1*. Also regionally confined were genes encoding fatty acid binding proteins 1 and 6 (*Fabp1* and *Fabp6)*, markers of domains A/B and E respectively, which encode different aspects of fat metabolism. In human tissue, domain A can be distinguished by human-specific domain A marker *ADA* and domain D by *PLB1*, as in mice. *SLC10A2* and *FABP6* are both expressed in domains D and E, with highest levels observed in domain E. These data support the patterns identified by scRNAseq and highlight the transitions in regional gene expression on a tissue level.

**Fig. 3.**
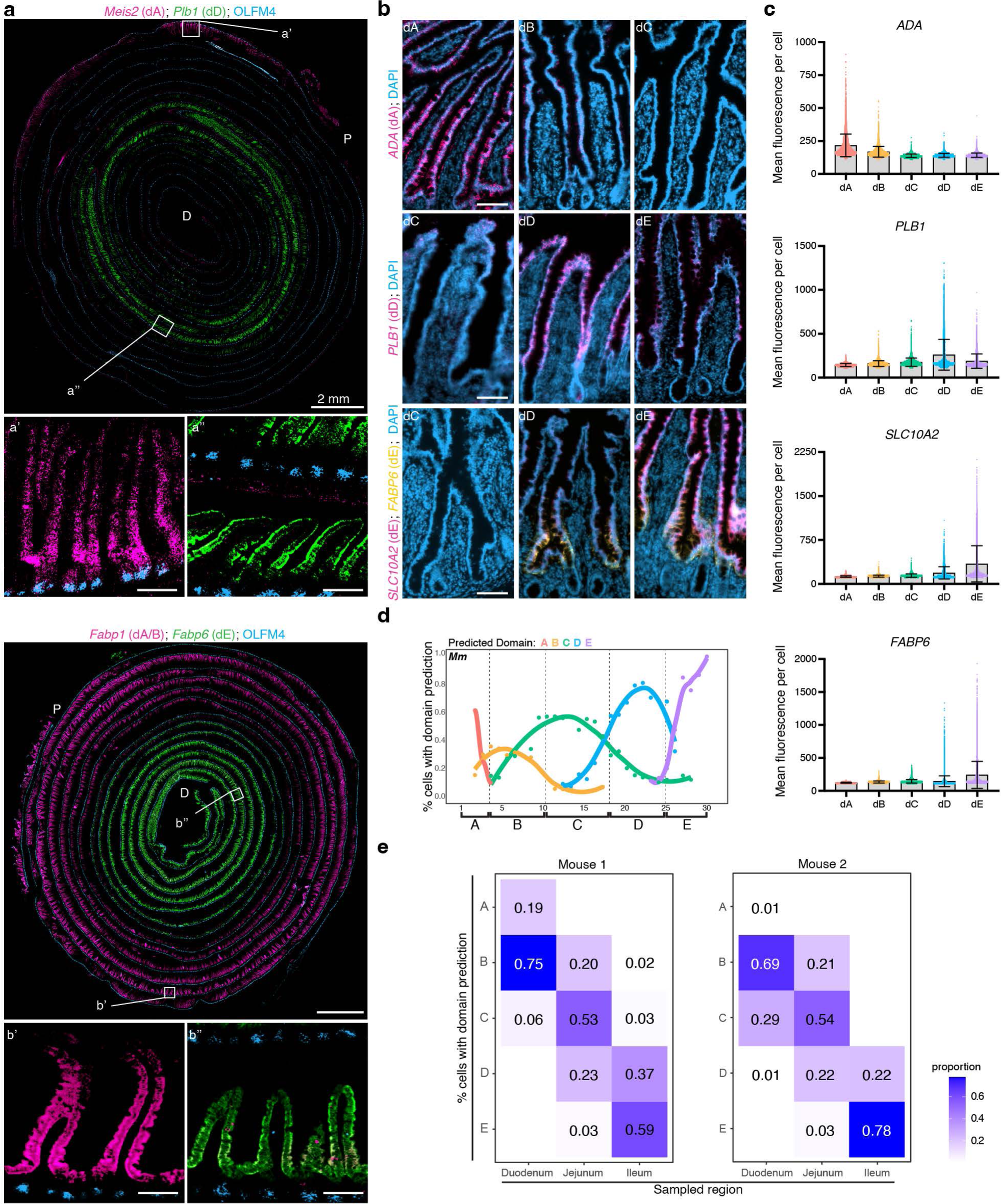
Domain identity can be detected across samples and used for systematic classification of intestinal regions. **a** Full-length murine intestinal tissue coiled from the proximal (outside) end to the distal (inside) end, probed with single-molecule ISH for select marker genes of domains as indicated. White boxes indicate insets. Scale bars are 2 mm, and 100 μm for insets. **b,c** images (**b**) of human tissue sections from indicated domains probed using single-molecule multiplexed ISH with indicated domain marker genes and quantification (**c**) of mean fluorescence per cell across 3-5 images per domain. Representative images and quantification from donor one are shown, n = 3 or 4 donors per domain. Scale bars are 100 μm. **d,e** Predicted domain identities of (**d**) enterocytes sequenced in mouse sequencing set two (test dataset, n = 2 mice) and (**e**) cells previously sequenced from two mice in published data^2^, as assigned by computational transfer of domain labels from the mouse dataset trained with known domain assignments (training dataset). In **d**, proportion of cells with the domain predictions at each segment position (x-axis) indicated by line color and dotted vertical lines indicate domain boundaries in training set in Fig. 1f,h. In **e**, proportion of cells in the reported classic intestinal regions are as indicated in each column. d Domain, Mm mouse.

We then sought to use the domain structure we defined in our mouse data to predict domains in other datasets. We employed a machine learning approach called transfer learning^18^ to train a classifier on the gene expression patterns of the domains defined by our mouse data. We then used the classifier to predict domain identities of enterocytes from a second cohort of two mice for which we collected data from 30 segments using the same procedure as in Fig. 1A (Extended Data Fig. 12). The domain boundaries inferred from the predictions were largely consistent with the boundaries defined in our first cohort (only the boundary between domains B–C was shifted by 2-3 segments, Fig. 3d). These data indicate that the discrete nature of the five domains can be used to predict the domain positions in other datasets.

We then used the trained classifier to predict the domain identities of cells sequenced in the original single-cell survey of the murine small intestine^2^, in which cells were categorized as deriving from the duodenum, jejunum, or ileum (Fig. 2e). Without a consistent method to define regionality within the intestine, we could not align our domain assignments to these traditional regions with precision, but based on the authors’ methodologies we estimated that the duodenum would align predominantly with domains A and B, the jejunum with B–D, and the ileum with E and a small portion of D. We found that domain predictions for most or all cells deriving from the duodenum, jejunum, and ileum aligned closely with our expectations. In the second sample sequenced in the original study, the model predicts fewer domain A cells, and more domain C cells, in the duodenal sample than expected, which may reflect minor differences in sampling strategies and is consistent with our observation that the position of the domains B–C boundary is more difficult to predict than others (Fig. 3d). Overall, the machine learning results support the presence of multiple distinct and recognizable transcriptomic signatures that align with five domains in the small intestine.

### The five domains reflect distinct functional zones of nutrient metabolism

To broadly evaluate whether the five computationally defined domains reflect significant differences in intestinal function, we determined the metabolic activities of all differentially expressed genes in enterocytes from each domain and analyzed those associated with nutrient absorption (Fig. 4a and Supplementary Table 5). In both species, domains A and B were most strongly associated with metabolism of fatty acids; domain C with carbohydrate metabolism; domain D with chylomicron and lipoprotein metabolism (which was also highly enriched in domain C in human) as well as amino acid transport; and domain E with cholesterol and steroid metabolism. In line with the high degree of transcriptional overlap between domains A and B (Figs. 2c–f), these domains were associated with many common processes, although in mouse, domain A was uniquely associated with iron uptake, and in both species, it displayed distinct transcripts associated with ion handling. Although domain C was largely defined by lack of expression of genes found in other domains (Fig. 1f,g), it was characterized by the highest expression of genes belonging to the carbohydrate transcriptional program^19^, indicating that domain C also performs a distinct physiological role. We similarly analyzed relevant NMF components (Fig. 2e,f), which provided a more distinct view not restricted by domain boundaries, of the regional span of co-expressed genes that encode nutrient metabolism proteins (Extended Data Fig. 13). For example, the formation of chylomicrons was more significantly enriched in domains C and D as above but detected at lower levels across domains A–D in both species. Overall, the regional patterns we identified were highly similar between the mouse and human intestine and reflect major aspects of nutrient absorption.

**Fig. 4.**
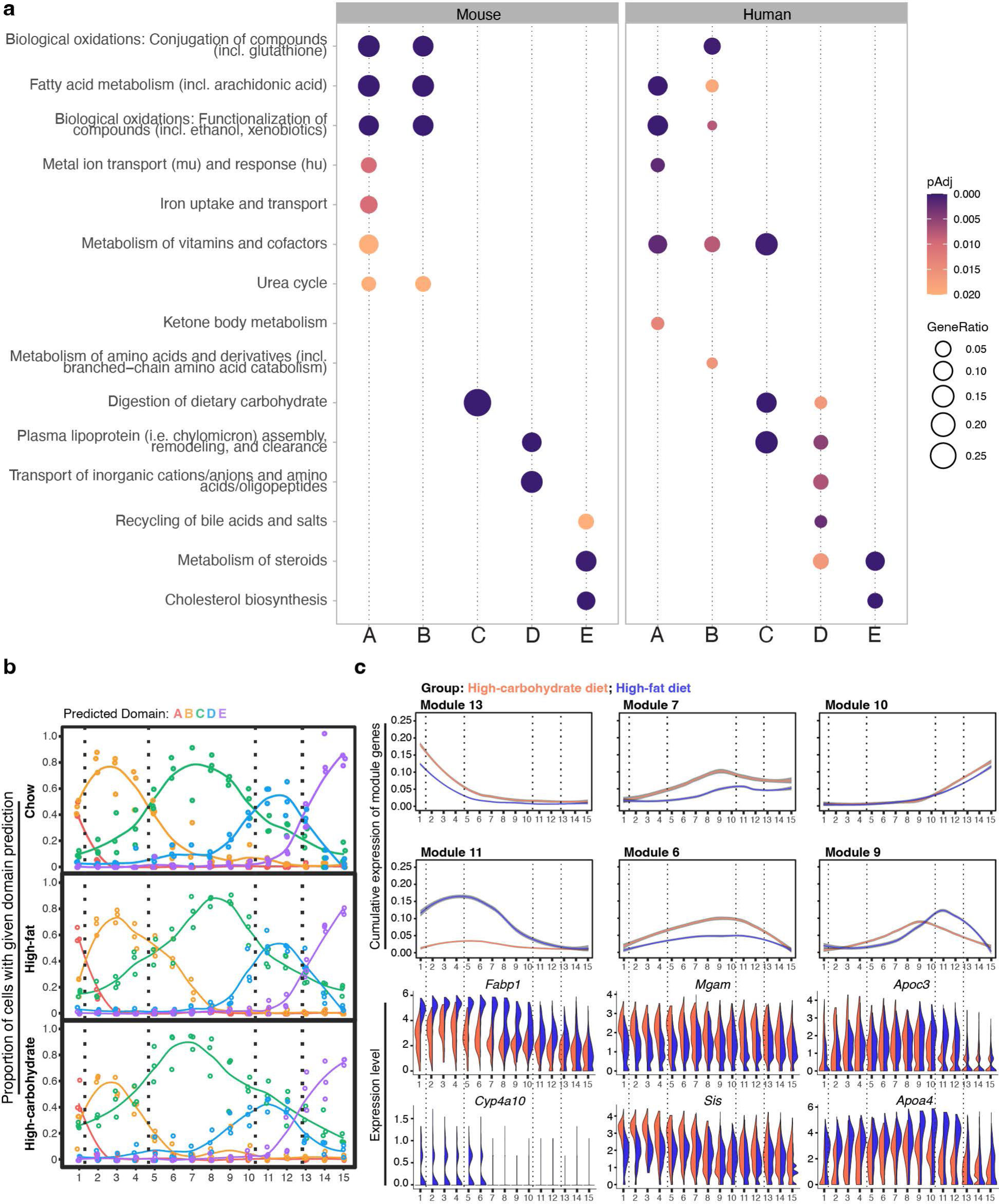
Domains are associated with distinct aspects of nutrient metabolism. **a** Summary of pathway enrichment in each mouse and human domain, represented as circles colored according to adjusted p-value and sized according to gene ratio (ratio of domain marker genes that are annotated with the pathway term). Selected domain-enriched, nutrient metabolism-associated pathways with adjusted p < 0.02 are shown. **b** Predicted domain identities of sequenced enterocytes from mice administered a high-fat or high-carbohydrate diet for 7 days (n = 3 mice per diet group), as assigned by computational transfer of domain labels from the mouse training dataset. Proportion of cells with the domain predictions in 3 mice per diet group indicated by color of best fit lines; dots indicate datapoints from each mouse. Dotted vertical lines indicate domain boundary positions predicted for chow diet group (top). **c** Cumulative expression of regionally variable NMF gene modules associated with nutrient metabolism across intestinal segments in each diet group, indicated by line color. **d** Expression level of select genes from the indicated modules associated with lipid metabolism (modules 11 and 9) and carbohydrate absorption (module 6) in mice fed high-fat (purple) or high-carbohydrate (orange) diets. Mm mouse, NMF non-negative matrix factorization.

These functional analyses suggest that the highest levels of lipid and carbohydrate metabolism occur in distinct domains when mice are fed standard chow: fatty acid metabolism most prominently in domains A and B, phospholipid metabolism in domain D, and carbohydrate absorption more broadly across the intestine but peaking in domain C. We hypothesized that enterocytes within these domains would differentially upregulate transcripts encoding the enzymes, receptors, and/or binding proteins needed to absorb an increased lipid or carbohydrate dietary load. To test this prediction, we fed mice either standard chow, a high-fat / low-carbohydrate diet, or an isocaloric high-carbohydrate / low-fat diet^19^. After 7 days (a time interval sufficient for enterocyte response to a change in dietary load^19-21^), we sequenced single epithelial cells as in Fig. 1a, this time from 15 equally sized segments across the intestine such that segment 1 corresponded to previously sequenced segments 1 and 2, and so forth. We obtained 27,881 high quality cells from the absorptive lineage (stem cells, TA cells, and enterocytes) from three mice for each diet (Extended Data Fig. 14). We applied the domain identity-trained classifier (Fig. 3d) to predict the domains of cells from mice fed each diet. The resulting prediction curves (Fig. 4b) were highly consistent across the three biological replicates per diet group and tracked the presence and position of five domains regardless of diet, in support of the robust nature of domain identity despite major dietary changes. Notable, however, was the broadening of the area associated with domain C in mice fed a high-carbohydrate diet into regions normally occupied by domains B and D (c.f. green line in segments < 5 and > 10 in high-carbohydrate diet, Fig. 4b). In segments of peak domain D prediction, a similar or higher percentage of cells were classified with a domain C identity, which may suggest that enterocytes with both domain properties co-reside at this position. This analysis suggests that enterocytes with domain C molecular and functional properties occupy a wider proportion of the small intestine, likely either in response to dietary lipid reduction or to carbohydrate augmentation.

We used NMF to examine gene modules and associated functions underlying this apparent shift in regional identity. Several, but not all, domain-associated modules were differentially expressed in mice fed high-fat versus high-carbohydrate diets (Fig. 4c, top half, Supplementary Table 4). Module 6 was strongly associated with carbohydrate absorption, and indeed we observed higher levels, over a larger region, of domain C signature genes that encode components of carbohydrate digestion, including maltase-glucoamylase (*Mgam*) and sucrase isomaltase (*Sis*), in mice fed a high-carbohydrate diet (Fig. 4c).

As previously noted, multiple NMF components collectively encode domain identity (Fig. 2e), and we also observed elevated expression of other modules such as 7 and 9 following high-carbohydrate feeding. Interestingly, module 9 included signatures of both domains C and D, and we observed a diet-selective response of genes within this component. Intestines from mice fed a high-fat diet upregulated domain D-associated module 9 genes as well as domains A and B-associated module 11, which were both functionally tied with lipid metabolism (Fig. 4c). Inspection of individual module components revealed that domain B genes known to play important roles in fatty acid metabolism^22^ and domain D genes in chylomicron assembly and triglyceride metabolism, were most strongly enriched, especially in their respective domains (Fig. 4d). Interestingly, domain E-associated module 10 appeared completely unaffected by these dietary interventions (Fig. 4c).

Together, hierarchical clustering of gene expression in single cells identified regionalized enterocyte domains in the mouse and human intestine that we experimentally validated using multiplexed ISH. Dietary challenge experiments demonstrated unique domain responses to individual nutrients and support the functional roles of domains A/B and D in lipid metabolism and domain C in carbohydrate absorption.

### Three regional stem cell populations reside within the small intestine

Having established patterns of specialized gene expression in enterocytes, we next asked at what stage of differentiation of the absorptive lineage these patterns emerge. As we captured a higher number of mouse than human stem cells per segment with scRNAseq, we focused this analysis on the murine absorptive lineage as a model. Theoretically, enterocytes could differentiate with little to no initial regional identity and take on local metabolic programs in response to microenvironmental cues; alternatively, enterocyte fate could be pre-determined by regionalized subpopulations of stem/progenitor cells. We found that mouse ISCs displayed localized gene expression (Fig. 1d), although less markedly than enterocytes, with 46% of genes expressed by crypt cells significantly varying along the proximal-distal axis (q < 0.05 for genes with mean sum-normalized expression above 5 X 10^-6^). We again applied Euclidian (Fig. 5a) and Jensen-Shannon (Extended Data Fig. 15a) distance metrics to calculate expression distance and perform hierarchical clustering of ISCs based on the top 100 most regionalized genes in this cell type. Hierarchical clustering showed that murine ISCs assembled into 3 regions well supported by the gap statistic (Fig. 2b) and with boundaries that fell within 2 segments of each of those that delineated absorptive domains B/C and D/E. JSD also indicated three groups, albeit with slightly different boundary positions. We favored the positions established with Euclidian distances as they draw directly from the gene expression matrix rather than a 2D projection. We refer to these populations as regional ISCs 1–3.

**Fig. 5.**
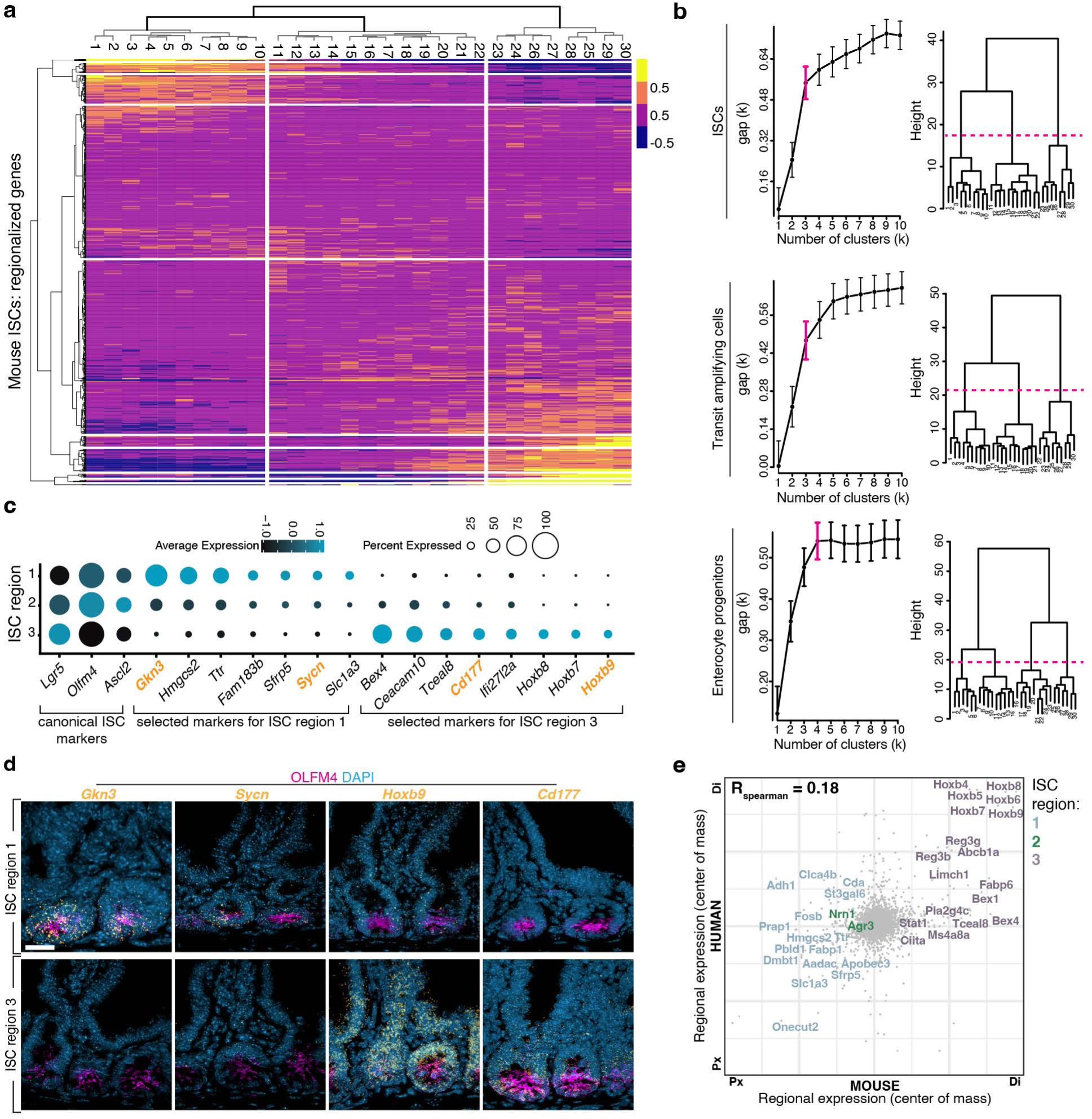
Three regional stem cell populations reside within the small intestine. **a** Average expression of the top 100 upregulated genes in murine ISCs in each segment, with segment order and hierarchical clustering based on expression distance between segments. Vertical white lines show the three domains that divide the ISC compartment, based on gap statistics. **b** Left: gap statistics for clusters of regional gene expression in regional ISCs, transit amplifying cells, and enterocyte progenitors. Right: cuts of dendrograms (dotted magenta lines) with optimal cluster numbers (magenta brackets, left) for each cell type. **c** Selected regional ISC subpopulation marker genes represented as dots colored according to average expression level and sized according to percent of ISCs expressing the marker. Bold orange marker labels were validated with ISH (Fig. 5d). **d** Intestinal crypts probed with single-molecule ISH for select regional ISC marker genes as indicated. Scale bars are 20 μm. ISCs intestinal stem cells. **e** Comparison of segment centers of mass for 7,668 homologous genes in mouse and human crypt cells with mean sum-normalized levels >1x10-5 in at least one point along intestinal length in both species. RSpearman = 0.18, p = 6.74 x 10-55, n = 2 mice and 2 human donors. Top segmentally variable genes in each species are shown, of which mouse regional ISC signature genes are color-coded as indicated. Px and Di identify the proximal and distal ends of the mouse (x-axis) and human (y-axis) small intestine.

As ISCs constitute only ∼1% of the total intestinal epithelium, they have been minimally sampled in previous reports, and our progenitor enrichment strategy enabled detection of new regional ISC markers (Supplementary Table 6). For example, in addition to known proximal and distal ISC markers (e.g., *Gkn3* and *Aadac* in region 1 and *Bex4* in region 3^2,23^), ISCs differentially expressed *Ttr* and *Sycn* in region 1 and *Cd177* in region 3 (Fig. 5c). In line with previous reports^23,24^, we observed bacterial response genes *Defa21 and Defa22* enriched in region 3 ISCs (Supplementary Table 6), suggesting a possible role for the regional microbiome or immune environment in shaping crypt zones.

We confirmed the spatial specificity of a subset of ISC markers using single-molecule ISH (Fig. 5d and Extended Data Fig. 15c–f, 16a,b). Whereas many markers were exclusively expressed by early-lineage cells (Extended Data Fig. 15b), we also noted a few shared regional markers between ISCs and later lineage cells such as hydroxymethylglutaryl (HMG)-CoA synthase 2 (*Hmgcs2),* which encodes a ketone body production enzyme. Expression of *Hmgcs2* expanded dramatically across the small intestine in response to a fat free diet, as would be expected upon initiation of ketogenesis, but other regional ISC markers such as *Gkn3* and *Bex1* remained stable regardless of dietary lipid levels (Extended data 15g). Furthermore, although regional gene expression in mouse and human crypt cells was not as tightly correlated as for enterocytes (Fig. 5e, RSpearman = 0.18, p=6.74e-55), many transcripts such as the classic regional identity marker *Onecut2* in region 1 ISCs, and Hoxb genes and *Bex1* and *4* in region 3 ISCs, showed similar expression profiles in both species.

We then used hierarchical clustering to model the point in the absorptive lineage at which these groups branch into 5 distinct enterocyte domains. We calculated the average expression of the most highly regionalized genes in TA cells and enterocyte progenitor cells from each segment, performed hierarchical clustering on the resulting data, and used the gap statistic to determine the optimal number clusters formed by these cell types. Our analysis indicated that 3 stem cell populations give rise to 3 TA cell populations, which then give rise to 4 groups of enterocyte progenitors that ultimately specialize into 5 distinct enterocyte populations (Fig. 1h and 5b).

### Transcriptional control of enterocyte regional identity

Given the broad zonation detected in early absorptive lineage cells (Fig. 5b, ISCs and TA cells), we wondered whether regionalized programs in ISCs might contribute to establishing the fate of enterocytes in each domain. In line with this possibility, previous reports^12-14^ have demonstrated that regional gene expression is maintained through long-term culture of organoids, and we observed maintenance of domain signature genes (Supplementary Table 2), including 27% of domain A genes and 30% of domain E genes, in their respective domain-specific organoid cultures (Fig. 6a, > 2.0 fold change, < 0.1 FDR, and qPCR validation of select signature genes in Extended Data Fig. 16c). While mesenchymal Wnt signals drive anterior-posterior small intestinal patterning during morphogenesis^25,26^, retention of location-specific transcript levels *in vitro* suggests that in the adult organ, some aspects of regional specialization are encoded within epithelial cells. Indeed, the best known small intestinal patterning factors, PDX1 and GATA4^26-31^, are expressed by epithelial cells.

**Fig. 6.**
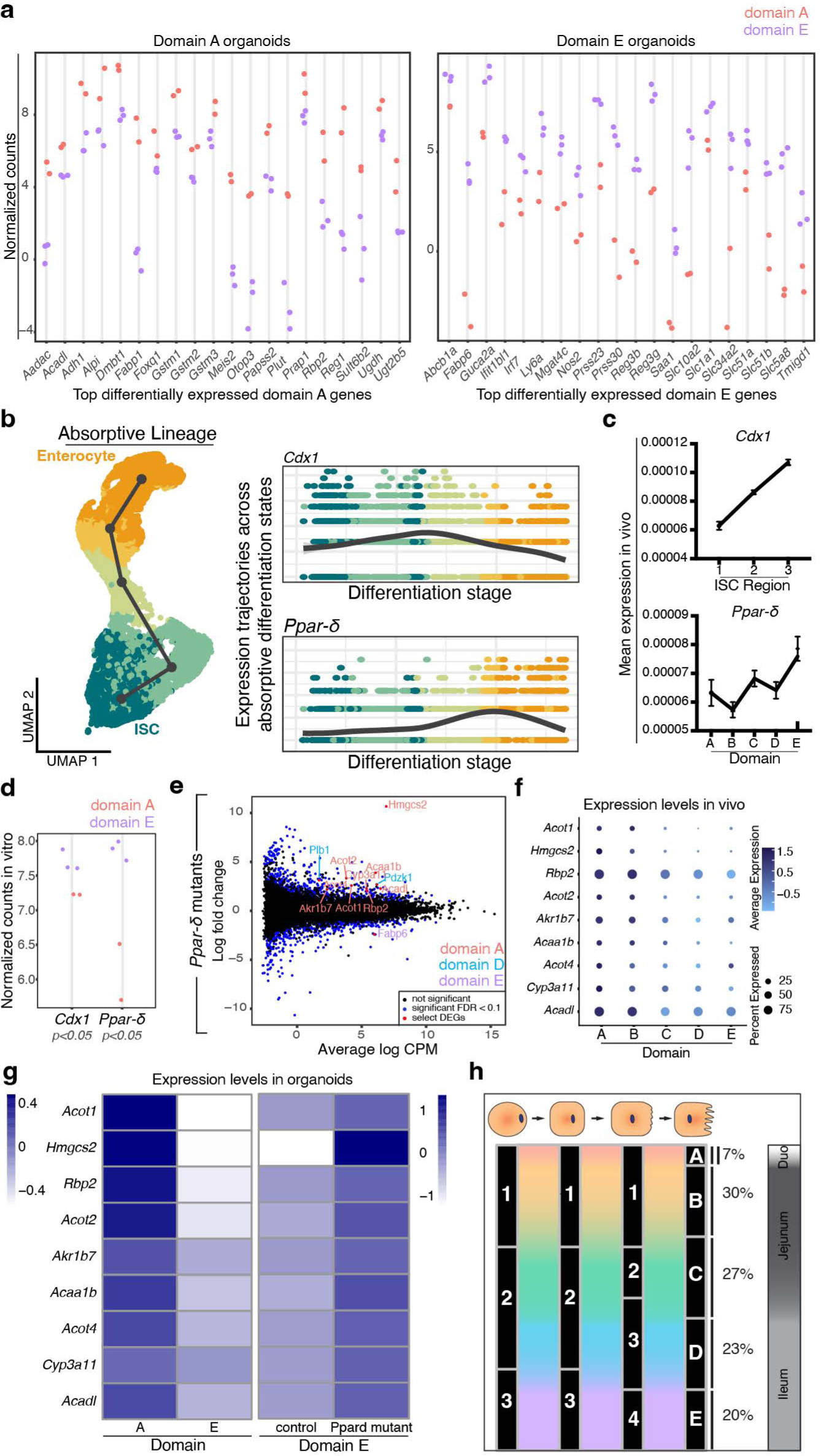
Transcriptional control of enterocyte regional identity. **a** mRNA levels of the top 20 domain A (left) and domain E (right) signature genes most highly differentially expressed in domain A or E-derived organoids, respectively, 5–6 days after passaging in long-term (> 5 week) culture, evaluated with mRNAseq. n = 2 dA organoid lines and 3 dE organoid lines. **b** UMAP of all murine absorptive lineage cells (left) and expression trajectories of *Cdx1* and *Ppar-ẟ* (right), colored according to inferred differentiation stage. Transcription factor expression trajectories were plotted for cells in domain E. **c** Expression profiles of *Ppar-ẟ* in enterocytes across domains and *Cdx1* in crypts across ISC regions. Data are mean expression levels of cells in indicated positions from mouse scRNAseq data, +/- standard errors of means, q < 0.01 for both genes. **d** mRNA levels of *Cdx1* and *Ppar-ẟ in* domain A or E-derived organoids, as in a. **e** Mean-difference plots of expression in *Ppar-ẟ* mutant organoids relative to controls. Dot colors specified in key. Regionally variable DEGs that encode lipid metabolism are labeled and colored by domain as indicated. n = 3 unique *Ppar-ẟ* mutant organoid lines and 2 control lines. **f** Dotplot of *in vivo* expression levels (analyzed in scRNAseq data) of identified DEGs in *Ppar-ẟ* knock out organoids. Dot size represents percent expressing enterocytes, color intensity represents average expression levels. **g** Heatmap showing mRNA levels of domain A lipid metabolism signature in domain A- and E-derived organoids as in a, and in control and *Ppar-ẟ* knock out domain E organoids as in e. **h** Summary of conclusions and model for regional specialization of the small intestine. Within the absorptive lineage (schematized, top), we find that ISCs occur in 3 regional populations, which likely give rise to 3 transit amplifying cell populations, which produce 4 enterocyte progenitors that ultimately specialize into 5 distinct mature enterocyte types that occupy absorption domains A–E. The estimated proportion of intestinal length of each domain and our approximation of corresponding traditional intestinal regions are shown.

To advance our understanding of the mechanisms that delineate the small intestinal domains defined here, we generated a model of epithelial-intrinsic transcription factors predicted to control the identity of every domain. We first used the gene regulatory network inference tools ChEA3^32^ and SCENIC^33^ to construct, from scRNAseq data, a predictive model of the transcription factors that are most likely to control domain-specific gene expression in enterocytes (Extended Data Fig. 17, Supplementary Tables 7 and 8). Notably, highly ranked factors on our list included established the zonation factors *Pdx1*^26-31^ and *Gata4*^26-31^, but many others were factors not previously associated with zonation.

Domain E is delineated from domain D by a sharp transition in expression of *Fabp6* and other domain-specific genes (Fig 2b), and it appears to be disproportionately affected by several largely regionally confined gastrointestinal diseases such as ileitis and necrotizing enterocolitis. Thus, we focused on domain E as a test case. We first ordered all enterocyte lineage cells in the domain according to inferred differentiation stage using slingshot^34^, allowing us to plot expression of each putative patterning factor across differentiation states (Extended Data Fig. 18a). Factors generally showed one of two trajectory patterns: highest expression in early lineage cells that declines as enterocytes differentiate, and expression in differentiated enterocytes or their immediate progenitors rather than early lineage cells(Extended Data Fig. 18b,c).

As we hypothesized that domain identity in enterocytes might be controlled at the level of ISCs, we first focused on putative patterning factors expressed most highly by ISCs and TA cells. Prominent among these candidates were homeobox genes that pattern the early gastrointestinal tract, but whose role in pattern maintenance during adulthood is less well understood. *Caudal type homeobox1 (Cdx1)* was expressed most highly in early-lineage cells (Fig. 6b) and specifically in region 3 ISCs and distal human ISCs (Fig. 6c and Extended Data Fig. 18d)*. Cdx1, 2*, and *4* are important regulators of hindgut patterning^35^. While the importance of *Cdx2* for the structure, function, and gene expression of the adult intestine is clear^36,37^, the role of *Cdx1* in the adult intestine has been more challenging to determine^36,38^.

To test our prediction that *Cdx1* maintains the metabolic profile of distal regions during adult homeostasis, we used two CRISPR-Cas9 gene editing strategies (Extended Data Fig. 19a,b, resulting in two batches of expression data) to delete the gene in domain E organoids, in which its expression is normally elevated relative to domain A organoids (Fig. 6e). *Cdx1* mutant organoids showed a trend towards decreased expression of the predicted target gene *Fabp6* that was consistent in both batches (Extended Data Fig. 19c,d); *Fabp6* is a domain E marker that is stably maintained in domain E organoids (Fig. 6a and Extended Data Fig. 16c). These data support our prediction that *Cdx1* promotes expression of the principal gene controlling long-chain fatty acid metabolism in the distal intestine, and more broadly, that regional patterning factors expressed as early as the ISC stage can control downstream aspects of nutrient processing and domain identity in enterocytes. It is likely that other patterning factors in the small intestine, such as *Gata4*, which is known to repress expression of several distal genes including *Fabp6,* function in concert with *Cdx1* to control domain E identity^28^.

We also tested our prediction that *Ppar-ẟ*, a known regulator of fatty acid oxidation and intestinal metabolism^39-41^, controls enterocyte genes associated with lipid processing in domain E. *Ppar-ẟ* modulates ISC metabolic response to diet^40,41^, and while we observed expression in early-lineage cells, this transcription factor was representative of those enriched in late lineage cells (Fig. 6b). *Ppar-ẟ* was expressed at slightly higher levels in domain E than in other domains in mouse and human (Fig. 6c and Extended Data Fig. 18d), a pattern that was recapitulated in long-term organoid culture (Fig. 6d). We performed CRISPR-modified deletion of *Ppar-ẟ* in domain E organoids in the same manner as described for *Cdx1*.

Bulk RNAseq of *Ppar-ẟ* mutants and controls, and qPCR validation of a subset of results, revealed differential expression of genes and enriched pathways associated with fat metabolism, including known PPAR target genes (Fig. 6e and Extended Data Fig. 19c,e,f). We observed decreased expression of domain E marker *Fabp6* and increased domain D-associated phospholipase (*Plb1*) levels. Interestingly, we observed upregulation of several genes that encode fatty acid metabolism enzymes such as ACADL, and ACOT1 and 4, that are specifically expressed in domain A in vivo during homeostasis (Fig. 6f), that are maintained in domain A organoid cultures (Fig. 6g). *Ppar-ẟ* loss in domain E organoids thus shifts regional organoids to a proximal lipid metabolism profile and supports our prediction that *Ppar-ẟ* maintains the expression signature of Domain E. *Ppar-ẟ* works in concert with proximally-enriched *Ppar-α*^41^; our results suggest that precise regional distribution of these factors may underlie PPAR-mediated patterning of lipid absorption across the intestine.

Collectively, these studies indicate that epithelial-intrinsic factors that are regionally expressed by cells at multiple stages of differentiation of the absorptive lineage participate in the stable maintenance of enterocyte domain identity across the adult intestine.

## Discussion

We have identified boundaries that divide the small intestine into five regional domains in both human and mouse, based on gene expression programs involved in nutrient absorption (Fig. 6h). Domain A, which likely represents the duodenum based on length and confined expression of the classic duodenal gene *Pancreatic and duodenal homeobox 1 (Pdx1)*, contained cells from segments upstream of the ampulla of Vater, where both bile and exocrine pancreatic secretions enter the intestine. A small set of domain A-specific genes rapidly declines in expression at the domain A-B boundary, including the homeobox gene *Meis2*, which represents a novel marker of this region; genes that encode subunits of the iron storage protein ferritin (*Fth1* in mouse, and, in a less starkly zonated manner, *FTL* in human); and genes involved in ion uptake.

Domain B overlaps with domain A in the first 6–10% of the intestine in both species; its proximal boundary is defined by termination of domain A-specific genes. Our analyses predict that these two domains are seeded by a common regional stem cell, and major physiological processes such as fatty acid metabolism occur in both domains. The gene constituents of neighboring domain C, which are most prominently associated with carbohydrate absorption, are also broadly expressed lengthwise, suggesting a wide range in which sugars are absorbed and metabolized. There are fewer positive markers of domain C than in neighboring domains, and we speculate that the presence of an intermediate region between domains B and D may allow for more plasticity to respond to environmentally induced shifts in transcriptional programs. In line with this possibility, domain C is the only domain that displayed a major size-wise change when mice were fed a reduced fat / increased carbohydrate diet. Further, the hierarchical clustering approach defines domain C in the second human donor more narrowly than in the first donor and in the mouse, possibly due to dietary differences. However, we also note that the expression of the domain C-defining NMF module in the second donor is broader than the hierarchical clustering results suggest. We believe that this difference reflects our overall conclusion that the boundary between domains B and C is not sharply defined and is subject to changes in response to environmental stimuli, but that these domains are delineated by independent molecular profiles that encode proteins required to execute non-overlapping functions.

Genes that encode ileal-specific functions, such as vitamin B12 uptake (*Cubn*) and bile salt recycling (*Slc10a2* and *Fabp6*), are enriched in domains D and E, suggesting that these regions best approximate the ileum, although our classification of previously published data suggest that domain D is likely included in studies of the murine jejunum. In both mouse and human, domain D declines as domain E increases with a small degree of overlap between two distinct gene modules. The domain D associated module is responsible for amino acid uptake and plasma lipoprotein processing and, as demonstrated by our dietary lipid modulation studies, is highly responsive to changes in dietary lipid loads. Domain E is predicted to function instead in metabolizing steroids and cholesterol, and remarkably, was found in our studies to be perfectly stable alongside substantial remodeling in the domain immediately adjacent in response to acute dietary change, suggesting that the intestinal area known as the ileum divides into two functional distinct parts. Future studies to evaluate whether this domain is innately less malleable, or whether it adapts to dietary cholesterol levels and cholesterol lowering drugs such as bile acid sequestrants, would be of significant interest.

The similarity of domain organization between mouse and human is striking, given the dietary and microbiome differences between humans and laboratory mice. Conservation and maintenance of spatial patterns of nutrient absorption across two mammalian species existing in radically different conditions supports the importance of an intrinsic intestinal positional system. *Ex vivo* maintenance of transcription factors including *Ppar-ẟ* and downstream target genes that define domain-associated metabolism lends further support to the idea that domain identity is hardwired in the adult intestine, presumably on a stem cell level. The three regional ISC populations identified here express factors predicted to direct specialization of enterocytes within the same regions, with *Cdx1* as one validated example by which *Fabp6* in enterocytes is controlled, at least in part, by a gene expressed most highly in stem cells. Several recent studies have demonstrated that metabolic programs such as ketogenesis^42^, fatty acid oxidation^43^, and sterol exposure^44^ can profoundly influence the fate decisions and regenerative behavior of ISCs and TA cells. These data add to our growing understanding of the roles of ISCs in defining local metabolic environments within the small intestine.

While core domain identities are stable, our studies demonstrate that gene expression levels and domain boundaries can adapt to nutritional cues. Further studies are needed to dissect the response of each domain to specific nutrients and other epithelial-extrinsic factors, such as the commensal microbiome and surrounding mesenchyme. Indeed, the small intestine has an impressive capacity to adapt to disruptions: bowel resection leads to a shift in expression of regional genes^45^, and parasite infection remodels crypt cell identity^46^, total intestinal length, and specialized cellular distribution^47^. How the epithelial-intrinsic organization and patterning mechanisms identified here may modulate and be modulated by the enteric microenvironment is an important question for future work.

A limitation of our study is human sample number; we sequenced the full-length intestines of two organ donors and performed selected validation for each domain on 2 additional donors. While we report salient aspects of domain organization across these individuals and species, analysis of additional subjects will strengthen our understanding of a core domain signature shared by humans and will undoubtedly reveal further intricacies that vary between people in diverse environments. A consequence of the number of human samples included, and of the greater variability between samples, is that our dataset was not sufficient to train a classifier to consistently recognize human cells in previously published datasets. For mice, however, we introduce a machine learning-based approach to identify the peak and boundary positions of five domains. This is the first systematic method to precisely track regions of the mouse intestine and provides a molecular classification system that future studies can utilize for consistent identification of relevant intestinal regions.

Finally, the similarities observed between mouse and human enteric regional organization have implications for understanding the regional distribution of gastrointestinal diseases that predominantly affect confined portions of the tissue, including celiac disease and adenocarcinomas in the proximal small intestine; and carcinoid tumors, lymphomas, necrotizing enterocolitis (NEC), and Crohn’s ileitis in the distal small intestine^48-50^. We note that NEC and ileitis most commonly affect domains D and E, which we found to be important sites of dietary fat response and metabolism, raising the intriguing possibility that lipid dynamics in these positions may modulate the local epithelial, immune, or microbial niche with relevance to these pathologies. This study provides a molecular roadmap that can be used to investigate the multifactorial interactions in specific cellular neighborhoods that may predispose specific regions to disease.

## Methods

## Mice

Male and female Lgr5^DTR-GFP51^ mice were used for scRNAseq and RNAscope experiments in Fig. 1 and Fig. 3, and female C57BL/6J (Jackson Laboratory Strain #000664, used 1 week after arrival) for diet modulation scRNAseq experiments.

Regional organoids to assess maintenance of regional signatures were generated from adult C57BL/6J mice; for CRISPR modulation from either Lgr5^creERT252^; *Rosa26^LSL-Cas9-eGFP/+^* (Jackson Laboratory strain #026175)^53^; ROSA26^tdTomato^ (Jax 007905)^54^ mice (strategy 1) or Lgr5^DTR-GFP51^ mice (strategy 2). Mice were 8–16 weeks of age at the start of each experiment. Previously defined^19^ specialized, purified high-fat / low-carbohydrate and high-carbohydrate / low-fat diets were purchased from Envigo and administered for 7 days. Rodent work was carried out in accordance with approved protocols by the Institutional Animal Care and Use Committee at the University of California San Francisco (UCSF).

## Human Intestinal Tissue

Human adult intestinal tissues were obtained from research-consented deceased organ donors at the time of organ acquisition for clinical transplantation through an IRB-approved research protocol with Donor Network West, the organ procurement organization for Northern California, in collaboration with the UCSF Viable Tissue Acquisition Lab (VITAL) Core. The first donor was a 44YO female with a BMI of 27 kg/m^2^ and the second donor a 30 YO male with a BMI of 25 kg/m^2^, both free of chronic and gastrointestinal diseases and cancer, and negative for hepatitis B/C, HIV, and COVID-19. Full-length intestinal tissues were collected after the clinical procurement process was completed, stored and transported in University of Wisconsin preservation media on ice, and delivered at the same time as organs for transplantation.

The study and all VITAL core studies are IRB-designated as non-human subjects research, as tissues are from de-identified deceased individuals without associated personal health information.

## Sample Dissociation

*Mouse Tissue:* Small intestinal tissues were removed from carcass and measured. The intestine from each mouse was lateralized, washed with RPMI (ThermoFisher) ‘FACS media’ supplemented with 3% FBS, Pen/Strep, Sodium Pyruvate, MEM non-essential amino acids, and L-glutamine, and cut into 30 pieces of equal length, or 15 pieces for dietary intervention studies. A single cell dissociation of the intestinal epithelium was obtained as previously described^19^. Briefly, tissue was incubated in the supplemented RPMI media described above with 5mM EDTA and 10mM DTT at 37°C with 5% CO2 for 20 minutes with agitation. Intestinal pieces were then triturated with a p1000 pipette, strained sequentially through 100 µm and 70 µm filters, and washed in RPMI containing 2 mM EDTA to separate the epithelial fraction.

### Human Tissue

Donated small intestines were stretched across an ice-covered trench drain and measured to be 546 cm (donor 1) and 667 cm (donor 2) long. These lengths were divided into 30 equal segments. 12 mm dermal punch biopsies (Acuderm inc.) and dissection scissors were used to collect 3–6 biopsies as technical replicates from within the central 4cm area in each segment. Punches were washed in DMEM/F12 (ThermoFisher) and PBS. Single epithelial cells were dissociated following previously published methods^55^. Briefly, cells were dissociated in Ca/Mg-free HBSS (ThermoFisher) with 10mM EDTA, Pen/Strep, HEPES, 2% FBS, and freshly supplemented with 5mM EDTA for 20–30 minutes at 37°C with 5% CO2 with agitation, and then for 15 minutes on ice. Cells were then triturated, treated sequentially with TrypLE (Gibco), DNAseI (Roche), and ACK lysis buffer as needed (ThermoFisher), and filtered through a 70 µm filter.

## Sample barcoding via MULTI-seq

Single murine and human cell suspensions from each segment were pelleted, washed, and resuspended with serum-free FACS media (as FBS and BSA prevent effective cell barcoding). MULTI-seq barcoding was performed as previously reported^15^: cells were suspended for 5 minutes on ice first with an anchor/barcode solution and then for 5 minutes on ice with a co-anchor solution. Following barcoding, cells from the proximal-most, middle, and distal-most 10 segments from mice and donor 1, and from segments with similar dissociated cell yields from donor 2, were pooled to help ensure relatively even sampling across the tissue length in subsequent steps.

## FACS

Pooled cells were stained with antibodies against CD45 (anti-mouse: BioLegend cat# 103130, anti-human: BD cat# 564047); EpCAM (anti-mouse: BioLegend cat# 118214, anti-human: BioLegend cat# 324208); and CD44 (anti-mouse/human: BioLegend cat #103026), and with DAPI. Live (DAPI–), single epithelial cells (CD45–, EpCAM+) with the exception of CD45+ tuft cells^2^, and progenitors (CD45–, Ep-CAM+, CD44+, Lgr5-DTR-GFP+ mouse cells and CD45–, Ep-CAM+, CD44+ human cells), were isolated using a BD FACSAria II equipped with FACSDiva Software Version 8 at the UCSF Parnassus Flow Cytometry Core. Plots were presented using FlowJo Version 10 (Extended Data Fig. 1).

## Single cell barcoding, library preparation, and sequencing

Sorted total epithelial and progenitor-enriched cells from each species were pooled separately before processing in individual lanes with the 10x Genomics Chromium system. Library preparation was conducted according to the 10x Genomics standard protocol, with modifications for MULTI-seq barcode library assembly as previously described^15^. Briefly, a MULTI-seq primer is added to the cDNA amplification mix. In the first SPRI bead clean-up step, the supernatant is transferred for a SPRI bead cleanup step. A PCR is also performed for MULTI-seq barcodes. Barcode libraries were analyzed using a Bioanalyzer High Sensitivity DNA system and sequenced.

Gene expression and barcode cDNA libraries were pooled and sequenced using an Illumina Novaseq 6000 machine at the UCSF Center for Advanced Technology (mouse samples and donor 2) and Institute for Human Genetics (donor 1).

### Analysis of single cell sequencing data

## Initial data processing

All analysis steps were performed using RStudio unless otherwise noted. Mouse set 1 sequencing reads were aligned using CellRanger version 3.0.1 (10x Genomics) to the mouse mm10-3.0.0 reference (10x Genomics). Sequencing reads for donor 1 were aligned using kallisto-bustools v0.46.2^56^ to the human GRCh38.95 reference. Sequencing reads for donor 2, mouse set 2 and the mouse diet experiment were aligned using CellRanger version 7.0.0 (10x Genomics) to the same respective references.

Raw gene expression count matrices were filtered using DropletUtils^57^ to identify real cells. Demultiplexing and removal of predicted doublets and unclassified cells was done with the deMULTIplex R package^15^ for mouse set 1 scRNAseq data; with the hashedDrops function of DropletUtils for donor 1 scRNAseq data; and with a combination of the hashedDrops function of DropletUtils and deMULTIplex2^58^ for the donor 2 scRNAseq, mouse set 2, and mouse diet data. Finally, identified cells were filtered according to number of UMIs per cell, number of genes per cell, and percentage of mitochondrial gene reads per cell (c.f. Extended Data Fig. 2, 4, 5, 12, and 14).

After performing sample demultiplexing on the murine set 1 and donor 1 scRNAseq data, we addressed two experimental issues computationally. First, in the murine scRNAseq data, we noted that identical MULTIseq sample barcodes were inadvertently applied to cells derived from segments 9–16 in the two mice sampled, as evidenced by the mix of male and female sex-linked genes in cells assigned to ‘Mouse A’, and a complete lack of cells in the same regions of cells assigned to ‘Mouse B’ (Extended Data Fig. 3). To distinguish between individual mouse samples, we used scPred^59^ to train a classifier that assigns cells from all segments to male, female, or unassigned status, and associated them to the appropriate segment position in mouse ‘A’ or ‘B’ accordingly (Extended Data Fig. 3b,c). Second, in the human scRNAseq data, we noted that human cells associated with the MULTIseq barcode for segment 30 were not recovered, which may be due to inefficient barcode labeling or sequestering of the barcode by dead cells or highly viscous mucus content in the distal-most portion of the human intestine during cell dissociation. All analysis of human data was therefore performed on segments 1–29, as displayed in the relevant Figures.

Mouse set 1 and donor 1 data were processed in Seurat V3^60^. Donor 2, mouse set 2 and the mouse diet experiment were processed in Seurat V4^60^. For mouse sets 1 and 2, total epithelial and progenitor-enriched samples were processed with the SCTransform function^61^ with 3000 features requested, with regression of differences in cell cycle state among cells, the level of expression of mitochondrial genes and of a set of sex-specific genes (Xist, Tsix, Ddx3y, Eif2s3y), followed by integration with Seurat’s IntegrateData function. Since the focus on mouse set 2 was on enterocytes, we did not integrate or further process cells from the progenitor-enriched fraction. The mouse diet samples were processed in the same way except for the regression of the expression of sex genes since all the mice in this dataset were females. Donor 1 total epithelial and progenitor-enriched samples were processed with the SCTransform function with 3000 features requested, with regression of the level of expression of mitochondrial genes, followed by integration with the fastMNN function. fastMNN integration was applied to the human scRNAseq data because it was the most effective procedure to correct batch effects between total epithelial and progenitor-enriched samples. Donor 2 total epithelial and progenitor-enriched samples were merged and processed with the SCTransform function with 3000 features requested, with regression of the level of expression of mitochondrial genes. Data from donor 2 did not require integration.

We performed data dimensionality reduction using principal component analysis in Seurat for all datasets except donor 1, for which the MNN components identified with fastMNN integration were used as low-dimension components. The number of principal components used was determined for each sample by inspection of the sample’s elbow plot. The following top components were used: mouse set 1, 50; mouse set 2, 32; mouse diet, 30; donor 2, 36; finally for donor 1 we used the first 50 MNN components. We also tested the stability of the downstream results (number of identified cluster, shape of the UMAP) to different choices of number of top principal components. Following dimensionality reduction, the nearest neighbor graph was calculated with the Seurat function FindNeighbors with the default argument k.param=20. We then identified clusters using the Seurat function FindClusters with default resolution (resolution=0.8), except for donor 1 for which we used a resolution of 0.55.

We classified the cell type identities of cells from mouse set 1 using Seurat to project previously reported reference cell type annotations for the murine intestinal epithelium^2^ onto the present data (Extended Data Fig. 2 and 6). Cell type annotation was refined by intersecting the transferred annotations and the clusters identified using Seurat, and resolving ambiguities using the following algorithm: ^57^ Clusters in which most cells had the same transferred annotation (this was the case for all clusters except cluster 15): cells annotated with the majority annotation were retained, cells without the majority annotation were annotated as “unknown” and not included in the analysis of regionality. (2) Cluster containing cells with two annotations transferred at high frequency: one cluster (cluster 15) contained mostly cells annotated as either “transit amplifying” or “enterocyte”. Cells annotated as one of these two types were retained, all other cells were annotated as “unknown” and not included in the analysis of regionality. Overall, cells of unknown identity constituted 7.6% of the total number of cell post-quality control in the mouse dataset but did not group into a single cluster.

All other single cells were annotated by assigning cell type identities based on marker gene expression^3^ (Extended Data Fig. 6, Fig. 12, Fig. 14). Clusters showing moderate expression of both cycling_g2m and enterocyte genes were annotated as “enterocyte progenitors; this annotation was also supported by the spatial observation that clusters annotated as enterocyte progenitors were found between TA cells and enterocytes in the UMAP visualization of the cells of the human dataset. Outlier cells that could not be annotated using existing marker genes (<2% of cells in either donor) were removed. Seurat was used throughout our analysis for the generation of violin plots, dot plots, ridge plots, and marker lists.

## Villus zonation scoring

Matlab version 2018b was used to annotate enterocytes according to their position along the crypt:villus axis using our previously published strategy^16^. Villus zonation scores draw from the summed expression of landmark genes^16^ and represent the ratio of the summed expression of the top landmark genes (*tLM*), and the summed expression of the bottom (*bLM*) and tLM genes (Extended Data Fig. 7). tLM and bLM were chosen based on the single cell-reconstructed zonation profiles as in^16^, as genes with a sum-normalized expression above 10^-3^ in at least one of the six villus zones and a center of mass above 3.5 for tLM or below 2.5 for bLM. The center of mass is average zone weighted by the expression of the respective gene^16^. An equal number of cells within the enterocyte clusters were assigned to each of 6 crypt:villus zones, Zones 1 – 6 (Extended Data Fig. 7).

## Calculation of % regionalization and gene expression distance across segments

The Kruskal-Wallis test was used to calculate the percent of regional zonation among genes with mean sum-normalized expression above 5 X 10^-6^. This analysis was only possible for cell types with > 40 cells per domain. Q-values were produced using the Benjamini-Hochberg procedure for multiple hypotheses correction. False discovery rate was set at q < 0.05. The centers of mass for all enterocyte-expressed genes (Fig. 2a), crypt-expressed genes (Fig. 5e), and gene markers of specific secretory cell types (Supplemental Table 1), were calculated across even fifths of the length of the intestine. For mouse-human correlations, we compared the segment centers of mass using a mouse-human orthology table based on Ensembl (version 109)^62^ using the BioMart data mining tool. Genes with a sum-normalized expression above 10^-5^ in at least one of the five segments are shown in the scatterplots in Fig. 2a and 5e. Genes with highest and lowest segmental centers of mass (reflecting proximal and distal-most expressed genes) and those with median centers of mass and highest Euclidean distance between the segmental profiles normalized to their maximum (reflecting center-most expressed genes) were labeled, and colored according to domain identity (Supplemental Table 2) if applicable.

Heatmaps were generated using pheatmap^63^ with the average normalized expression of the 150 genes most highly upregulated per segment in enterocytes (defined as the combination of cells annotated as differentiated or mature enterocytes) (Fig. 1f,g), or the top 100 marker genes per segment in intestinal stem cells (Fig. 5A). Because cell number per segment is variable in the human dataset, segments were grouped into pairs for this analysis. Heatmaps visualize data from a matrix in which each cell contains the average expression of a marker gene in each segment. Segments and genes were clustered based on the Euclidean distance between cells in the matrix. The optimal number of clusters was identified by computing the gap statistic using the clusGap function of the R package cluster (version 2.1.4) using default parameters. We also confirmed that domain divisions were stable when alternate numbers of top upregulated genes were used (Extended Data Fig. 8c, displaying 75–100 upregulated genes per segment).

To evaluate domain assignments with a different approach, we calculated the Jensen-Shannon Divergence (JSD)^64,65^ for enterocytes and intestinal stem cells on the mouse dataset (Extended Data Fig. 8d, 15a). To calculate JSD, we assigned a center of mass to each segment by bivariate Kernel Density Estimation and calculated pairwise JSD between the resulting vectors. For enterocytes, JSD was calculated for each mouse individually. Mouse 2, which contains less cells and has more cell number per segment variability than Mouse 1, had slightly weaker segment ordering (note the positions of segments 19-20) than Mouse 1, but mis-ordering was confined to domains and did not ultimately affect our interpretation of appropriate boundary divisions.

Domain-defining signature score (Fig. 2c,d) is a z-score metric representing the mean expression of the 20 most differentially expressed genes in a given absorption domain. The signature scores were computed from scaled and centered gene expression data following SCTransfom in Seurat.

## Non-negative matrix factorization analysis

We performed non-negative matrix factorization analysis using the cNMF package version 1.4 in R^66^. We used the raw count matrixes for a given subset of cells as input to cNMF, and ran cNMF with default parameters. For visualization of the results, we selected the 250 genes with the strongest contribution to a component and used the Seurat function AverageExpression to compute the averaged expression of the selected genes.

## Prediction of intestinal domain locations using transfer learning

We performed the computational transfer of domain labels from mouse datasets with known domain assignment (training datasets) to datasets with unknown domain position (test dataset) by transfer learning using the cFIT package version 0.0.0.90 in R^18^. We used the raw count matrixes for enterocytes and mature enterocytes as input to cFIT. All cells (both the training and test sets) were labeled according to their experimental batch. Cells from the training sets were also labeled according to their previously assigned domains. cFIT was run with default parameters and requesting 15 number of factors of the common factor matrix (shared across training and test datasets). We used the following datasets:

| TRAINING DATASET | TEST DATASET |
| --- | --- |
| Mouse set 1 | Dataset GSE92332_Regional_UMIcounts (GEO database) collected from duodenum, jejunum, and ileum<sup>2</sup> |
| Mouse set 1 | Mouse set 2 +<br>Mouse chow diet |
| Mouse set 1 + mouse set 2 +<br>mouse chow diet | Mouse fat-free diet +<br>Mouse high-fat diet |

## Functional pathway analysis

Pathways enriched in each mouse and human absorption domain (adjusted p value < 0.02, Fig. 4a, Supplementary Table 5) or regionally variable NMF component (adjusted p value < 0.04, Extended Data Fig. 13) were identified using the ReactomePA enrichPathway tool and compared using the clusterProfiler package^67^. Selected pathways associated with nutrient metabolism are shown. Pathways were edited to remove redundancy and plotted with ggplot2.

## Evaluation of transcriptional control of domain identity

We first used ChIP-X Enrichment Analysis 3 (ChEA3)^32^ to identify transcription factors predicted to control genes differentially expressed in enterocytes from each absorption domain. We repeated this analysis for enterocytes, TA cells, and ISCs, such that we might evaluate which transcription factors expressed by each of these cell types is predicted to control domain-specific expression in enterocytes. Transcription factor enrichment results generated with this approach (Supplementary Table 7) are based and ranked according to several types of data including transcription factor-gene association in RNAseq and ChIP-seq datasets, and co-occurrences in submitted gene lists. We also used SCENIC^33,68^ to infer Gene Regulatory Networks based on co-expression and motif analysis of transcription factors and targets which were then analyzed in individual differentiated and mature enterocytes (Supplementary Table 8). To evaluate expression of each transcription factor along stages of absorptive cell differentiation, from ISC to enterocyte, we used Slingshot^34^ to infer differentiation pseudotime for all absorptive cells and order the cells accordingly.

Transcription factors were evaluated according to their predictive rank in ChEA3, convergent identification in ChEA3 and SCENIC analyses, and regional expression across domains (Extended Data Fig. 17). We grouped transcription factors according to highest expression at early (ISC/TA cell) or late (enterocyte precursor or later) stages of the absorptive lineage (Extended Data Fig. 18).

### Visualization of regional marker transcripts

Full-length murine small intestinal tissue or transverse cross sections of human intestines from indicated domains were immersed in 4% PFA for 24-48h at room temperature and EtOH for 24 hours at 4°. Murine small intestines were coiled into a ‘swiss roll’ from an outer proximal tip to an inner distal tip. All tissue underwent standard dehydration and paraffin embedding.

The RNAscope Multiplex Fluorescent V2 Assay (Advanced Cell Diagnostics) was used according to the manufacturer’s instructions to probe for transcripts of interest. Entire swiss rolls were captured with a Leica DMi8 microscope equipped with LAS X Software and an automated stage, allowing for tilescan imaging of frames at a 20X magnification; 3-5 individual images were acquired per region from each donor. Regional patterns of selected individual marker transcripts were confirmed on at least three mice each and in 3-4 donors per domain, including the 2 donors sequenced in this study. Images of individual murine crypts and crypt-villus units were also captured using a Zeiss LSM900 confocal microscope.

For morphometric analysis of villus height (Extended Data Fig. 10), the lengths of tilescanned swiss rolls were tracked using a custom macro for Fiji^69^, allowing assignment of the precise positions of 30 equal segments. Villus base to tip distances were measured for 3-5 villi in each segment, for each of 4 mice. One-way ANOVA followed by Tukey’s multiple comparisons test for villus heights across all segments in each domain was performed using Prism software (GraphPad Prism version 8 for MacOS).

Human tissue images were analyzed using a custom script in QuPath software^70^. Briefly, nuclei detection was performed using StarDist2D and cell segmentation was performed with the cell expansion variable set to 10 µm. The mean fluorescence value for each cell was plotted (Fig. 3c), and one-way ANOVA to compare mean fluorescence in each donor by domain was performed (Extended Data Fig. 11b) using Prism.

### Investigation and genetic perturbation of regional organoids

## Generation and qPCR evaluation of regional organoids

Intestinal crypts were isolated from domains A-E of fresh intestinal tissue using methods previously described^71^.

For evaluation of gene expression with qPCR or mRNAseq, organoids that had been cultured for at least 1 month (5–13 weeks), and 5–6 days after passaging, were washed with PBS and resuspended in TRI reagent containing 1% 2-Mercaptoethanol. RNA was extracted using Direct-zol RNA Miniprep Plus (Zymo Research) and cDNA reverse transcribed with High Capacity cDNA Reverse Transcription Kit (Applied Biosystems) according to the manufacturer’s instructions. qPCR using the primers listed in Supplementary Table 8 was performed using a C1000 Touch Thermal Cycler (Biorad).

## CRISPR-mediated gene disruption

Two single guide RNAs (sgRNAs) were designed for each target using the Benchling CRISPR Guide RNA Design tool (https://www.benchling.com/crispr/). Following previously described methods^72^ and using BstXI (Thermo Fast Digest, cat: FD1024) and BlpI (Thermo Fast Digest isoschizomer Bpu1102I, cat: FD0094) restriction enzymes, we inserted a sgRNA into the pU6sgRNA-EF1alpha-puro-T2A-BFP single cassette vector, which expresses the mouse U6 (mU6) promoter and constant region 1 (cr1)^73^, and the second sgRNA into pMJ117, which expressed the modified human U6 (hU6) promoter and cr3^74^. sgRNA sequences, and primers used for subsequent PCR amplification (Q5 Hot Start High-Fidelity 2X Master Mix, NEB) of sgRNA expression cassettes are provided in Supplementary Table 8. pU6sgRNA-EF1alpha-puro-T2A-BFP was then digested with XhoI and XbaI (NE Biolabs) and gel purified along with PCR fragments. sgRNAs were then incorporated into the pU6sgRNA-EF1alpha-puro-T2A-BFP backbone using NEBuilder® HiFi DNA Assembly Master Mix (NE Biolabs) according to manufacturer’s instructions. Lentivirus was produced from the resulting dual sgRNA constructs by the UCSF Viracore. Virus was concentrated using Lenti-X Concentrator (Takara Biosciences).

To increase the efficiency of CRISPR mutagenesis, we also used a second strategy based on simultaneous delivery of Cas9 and sgRNA by lentiviral vectors. Using Esp3I restriction enzyme (New England Biolabs, cat: R0734S) we inserted each sgRNA into lentiCRISR v2 (Addgene 52961) which allows simultaneous expression of gRNA driven by U6 promoter and Cas9/PuroR driven by EF1alpha. Cloning was performed as described^75^, and successful insertion of sgRNA sequence was validated by Sanger Sequencing using primer 5’-GCACCGACTCGGTGCCAC-3’. sgRNA sequences are provided in Supplementary Table 9. Lentivirus was produced from the resulting vectors as described^75^.

Lentiviral transduction of adult, regional organoids for all experiments were performed as described^76^. Briefly, intestinal organoids were grown for at least 4 days prior to infection in “ENRWNTNIC” (50% growth medium/50% Wnt-cultured medium and 10mM nicotinamide), supplemented with 10uM Y-27632, and 2.5uM CHIR to induce spheroid formation. Stem cell-enriched spheroids were broken into single cells for the addition of viral mix containing 8ug/ml polybrene, followed by a 1 hour spinoculation, and a 6 hour incubation at 37°. Infected cells were then plated in Matrigel. Puromycin selection was performed 72 hours after recovery. Spheroids were converted to organoids over the course of approximately 7 days by gradual transition of ENRWNTNIC to ENR medium. Infected organoids were expanded and, for strategy 1, treated with 4-hydroxytamoxifen to induce Cre recombinase-dependent expression of Cas9 endonuclease and EGFP. From these cultures, organoids were passaged at a low density (strategy 2), or small numbers (1-100) of single, BFP+ (transduced), GFP+ (tamoxifen-induced) cells were sorted into individual, Matrigel-coated wells of a 96-well plate (strategy 1), in both cases allowing for precise manual isolation of individual organoids. After ∼10 days of growth, single mature organoids were collected and used for clonal expansion. To confirm genetic disruption, genomic DNA was isolated (Lysis and Neutralization Solutions for Blood, Sigma), genotyped with PCR, and the mutant alleles were sequenced (primers, Supplementary Table 9). Clones carrying the wild-type alleles were excluded and only the clones with deleterious alleles were used for the downstream analyses.

## mRNAseq of regional organoids

RNA was collected from confirmed mutant organoid clones, transduced organoids uninduced by OHT, and untreated organoids as described above for qPCR evaluation. All organoid lines were cultured for 5–6 days post-passaging to ensure consistent and complete differentiation status across samples. RNA sample QC, mRNAseq library preparation, and mRNAseq (Illumina, PE150, 20M Paired Reads) was performed by Novogene.

Genome indexing and quantification of transcript abundances by pseudoalignment were performed using Kallisto version 0.46.0^77^. Non-expressed genes were filtered by retaining genes with > 5 reads in at least 4 samples. RUVseq^78^ was used to control for “unwanted variation” between samples. Differentially expressed genes in mutant organoids compared to untreated organoids were identified using EdgeR. Since mutant organoids were assayed without replication, data dispersion was estimated from all but the 5,000 most variable genes in the entire dataset.

## Acknowledgements

We are grateful to Hikaru Miyazaki, David Castillo-Azofeifa, and other members of the Klein lab for valuable discussions, experimental assistance, and protocol development. We thank Michael Helmrath, Noah Shroyer, Yuan-Hung Lo and members of the Intestinal Stem Cell Consortium for critical scientific input throughout this project. Thank you also to Benjamin Ohlstein, Ina Chen, Keren Bahar Halpern, Zuri Sullivan, Danny Conrad, Jess Sheu-Gruttadauria, Jeffrey Bush, and Eric Chow for sharing data, resources, and expertise. This study benefited from the following cores and facilities at UCSF: the Center for Advanced Technology, the Institute for Human Genetics, Parnassus Flow Cytometry Core, ViraCore, VIable Tissue Acquisition Lab, and the Laboratory Animal Resource Center. Portions of schematic Figure panels were created with BioRender.com. This work was funded by NIH R35-DE026602 and by U01DK103147 from the Intestinal Stem Cell Consortium, a collaborative research project funded by the National Institute of Diabetes and Digestive and Kidney Diseases and the National Institute of Allergy and Infectious Diseases, to O.D.K. R.K.Z. was supported by NIH F32 DK125089 and an American Cancer Society – South Florida Research Council Postdoctoral Fellowship (PF-20-037-01-DDC). Finally, our most sincere gratitude to Donor Network West, and to the organ donor and the donor’s family for their generosity in supporting basic science research.

## **Data** Availability

The datasets generated and analyzed during the current study are available in the GEO repository, accession GSE201859.

## Author contributions

R.K.Z. and O.D.K. conceived and developed the study, R.K.Z. conceived and planned experiments, R.K.Z., C.S.M., S.I., and D.B. developed the analysis strategy and performed data analysis, D.B. conceived several computational approaches, and supervised and verified the analytical methods, R.K.Z. L.W., K.L.M., E.R., A.R., and V.N. carried out experiments, A.R.G. and J.M.G. facilitated the human intestinal tissue donation, Z.J.G., R.M.L, J.M.G, and S.I. provided intellectual review of project content, and R.K.Z., D.B. and O.D.K. wrote the manuscript with input from all authors.

---

**Extended Data Fig. 1.**
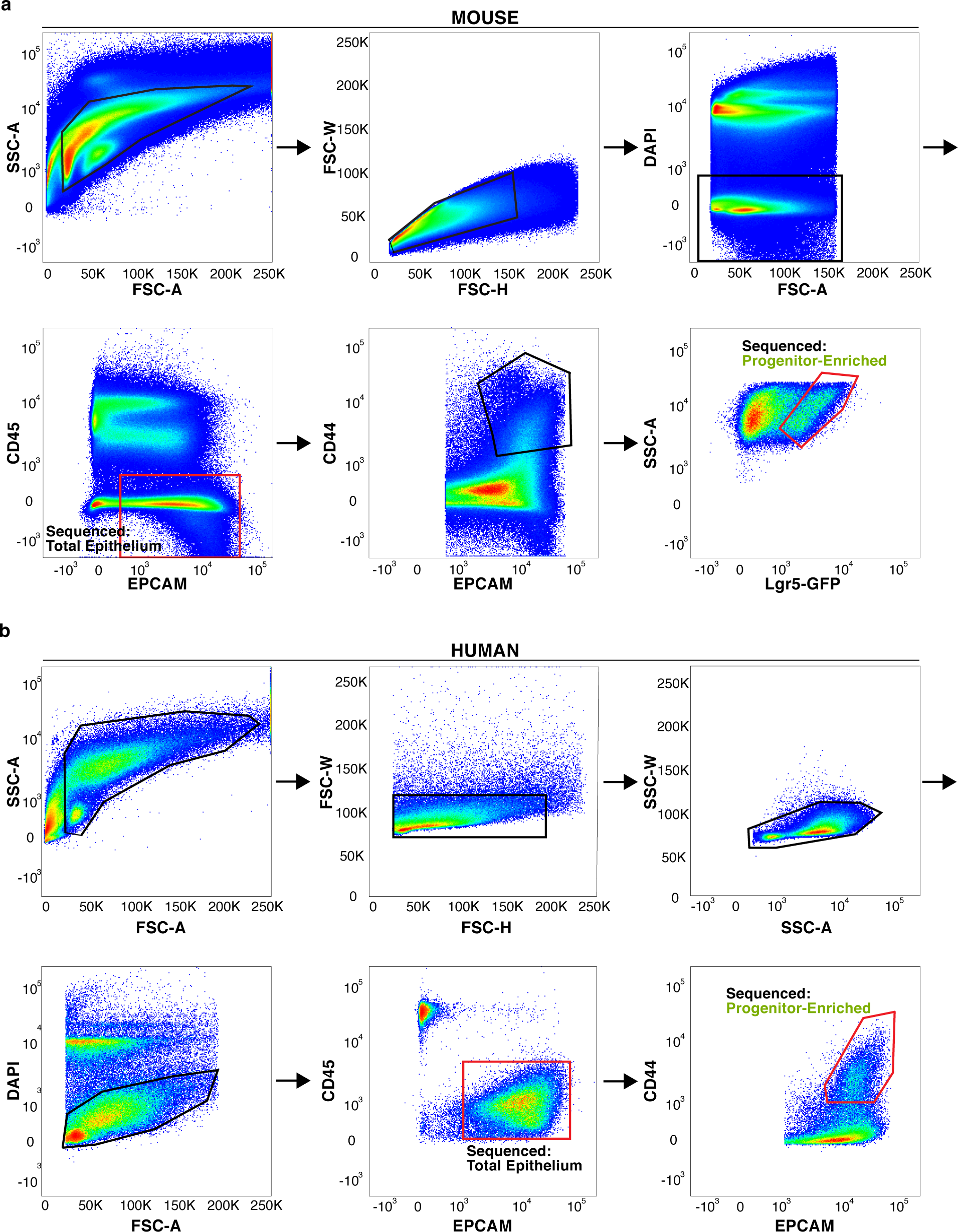
Strategy for isolation of murine and human epithelial cells for single cell RNA sequencing (scRNAseq). **a,b** Representative flow cytometry plots of sequential gating strategy for single, live (**a**) murine total epithelial (CD45–, EPCAM+) and progenitor-enriched (CD45–, EPCAM++, CD44++, Lgr5-GFP+) cells and (**b**) human total epithelial (CD45–, EPCAM+) and progenitor-enriched (CD45–, EPCAM+, CD44+) cells. CD45+ tuft cells were not captured in this study. FSC, forward scatter; SSC, side scatter.

**Extended Data Fig. 2.**
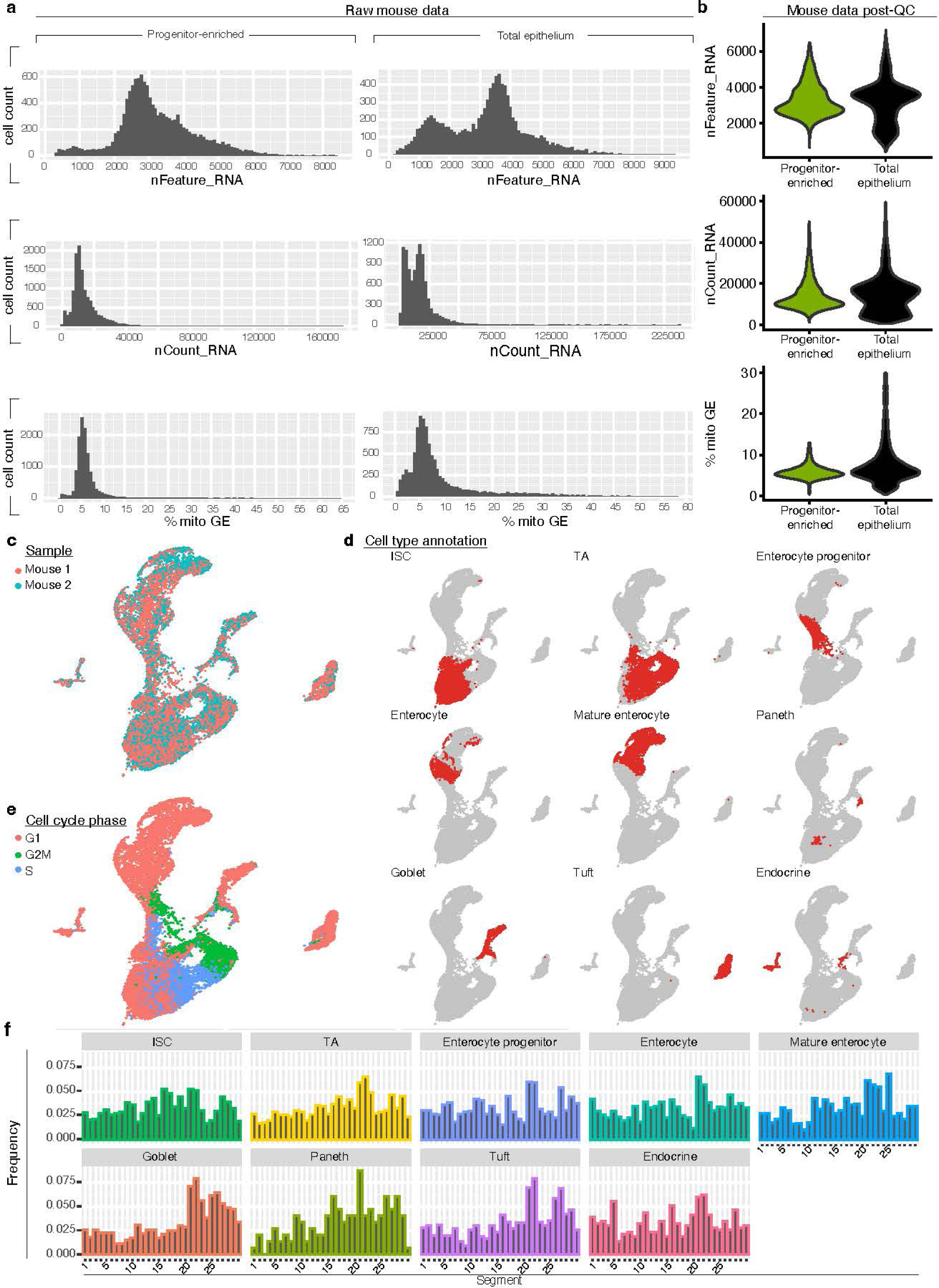
Quality control and initial processing of mouse scRNAseq data. **a,b** Quality control metrics of data, including number of genes detected (‘nFeature_RNA’), number of unique molecular identifiers detected (‘nCount_RNA’), and percent mitochondrial reads (‘% mito GE) before (a) and after (b) processing data. **c-e** Uniform Manifold Approximation and Projection for Dimension Reduction (UMAP) of total murine epithelial cells sequenced post-QC, colored according to mouse identity (**c**), cell type annotation (**d**), or cell cycle phase (**e**). **f** Frequency of epithelial cells of indicated subtype by segment. QC, quality control, mito, mitochondrial; GE, gene expression; ISC, intestinal stem cell; TA, transit amplifying; G1, growth 1; G2M, growth 2 mitosis; S, synthesis.

**Extended Data Fig. 3.**
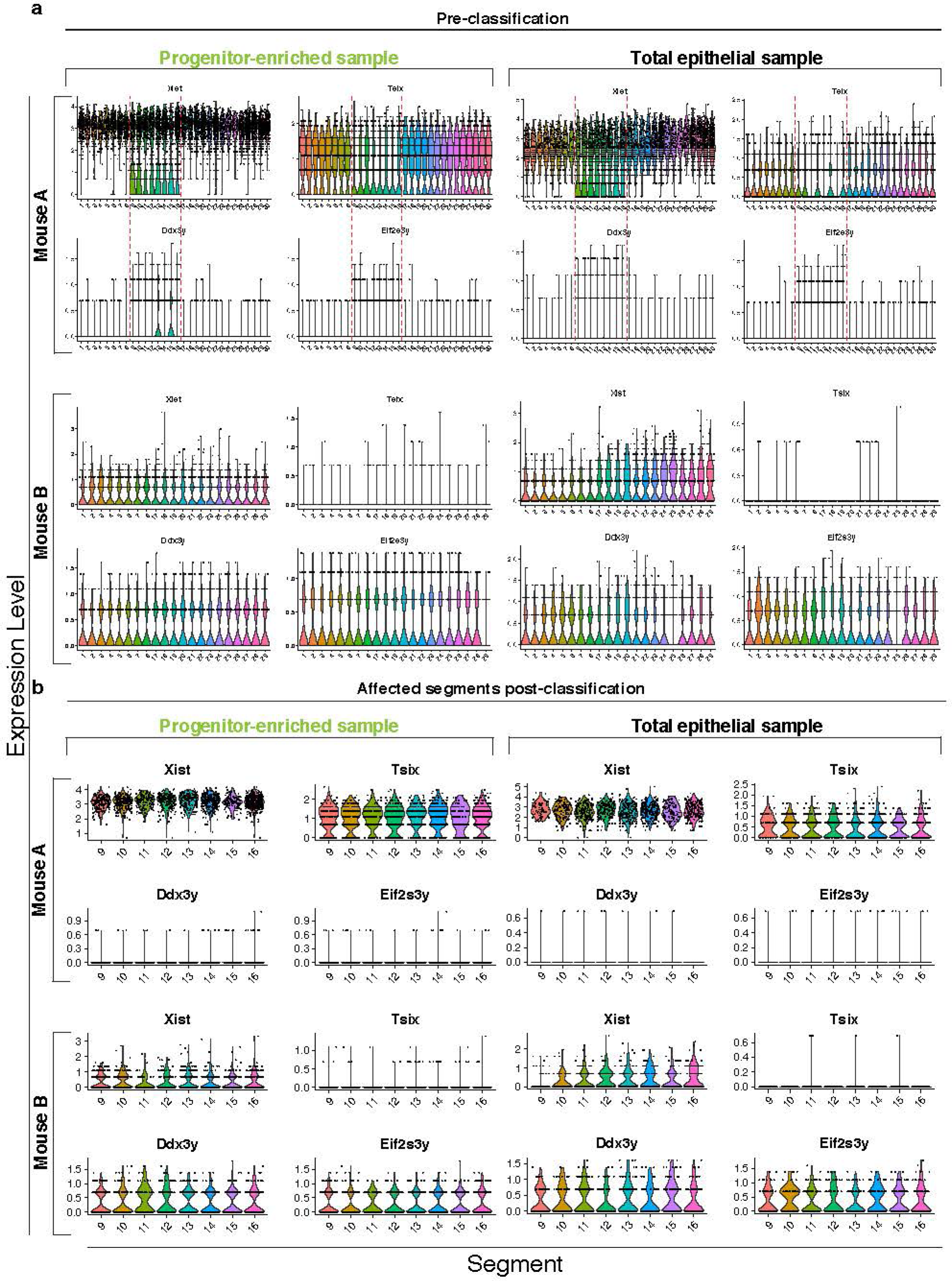
Classification of murine cells with mouse identity. **a** Expression of sex-linked genes in progenitor-enriched (left) and total epithelial (right) cells from each mouse prior to classification. A mix of male and female-linked genes were evident in segments 9-16. **b** Expression of sex-linked genes in progenitor-enriched (left) and total epithelial murine (right) cells from each mouse after training classifier to assign cells from all segments to male, female, or unassigned, and associate them with the appropriate segment positions in mouse ‘A’ or ‘B’. Classification and reassignment of cells resulted in exclusive expression of either female or male-linked genes in Mouse A and Mouse B, respectively.

**Extended Data Fig. 4.**
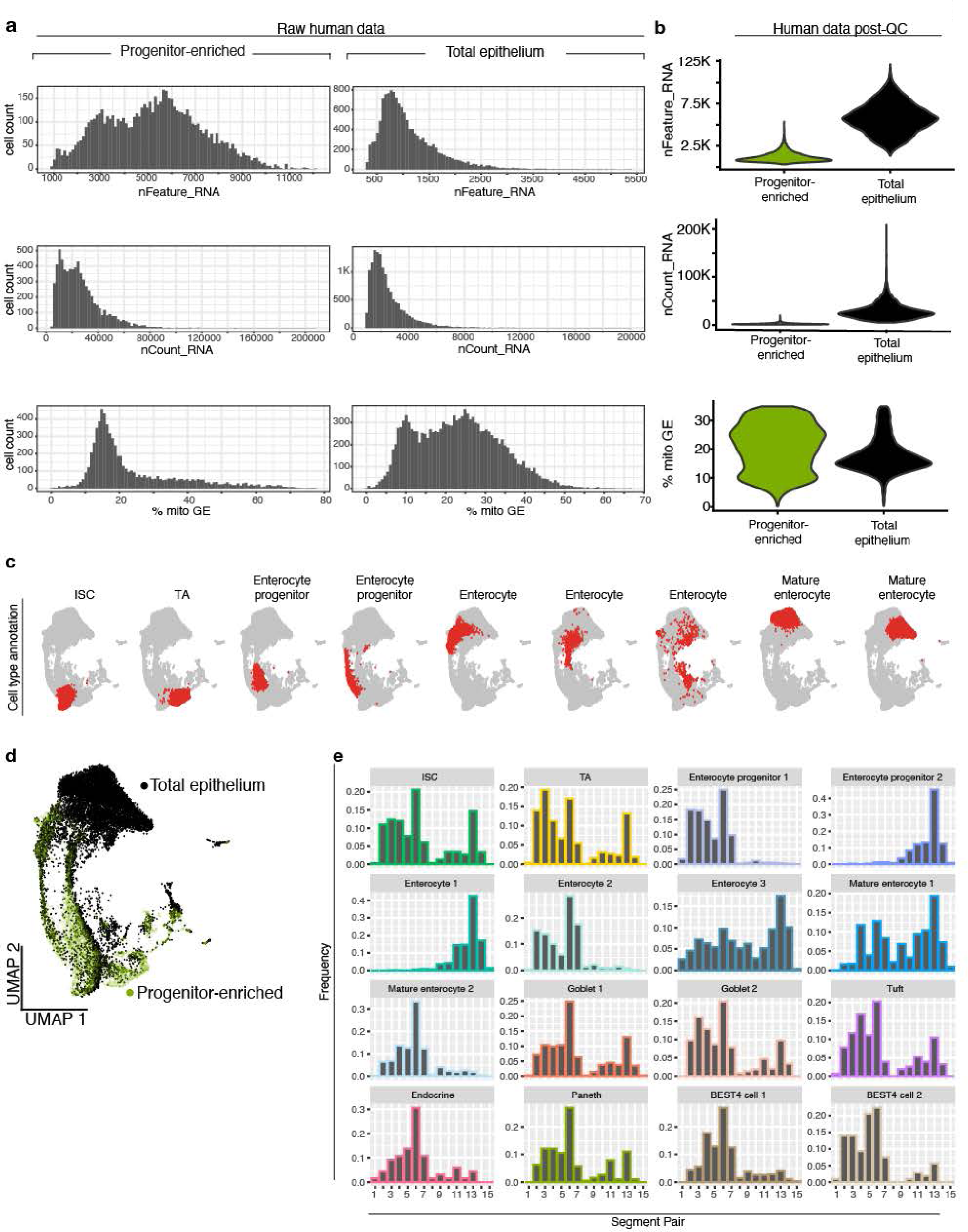
Quality control and initial processing of human scRNAseq data from human subject 1. **a,b** Quality control metrics of data, including number of genes detected (‘nFeature_RNA’), number of unique molecular identifiers detected (‘nCount_RNA’), and percent mitochondrial reads (‘% mito GE’) before (**a**) and after (**b**) processing data. **c,d** Uniform Manifold Approximation and Projection for Dimension Reduction (UMAP) of total human cells sequenced post-QC, highlighting cell type annotation (**c**) and total epithelial or progenitor-enriched sample identification (**d**). **e** Frequency of cells of all epithelial subtypes by segment pair. QC, quality control, mito, mitochondrial; GE, gene expression; ISC, intestinal stem cell; TA, transit amplifying.

**Extended Data Fig. 5.**
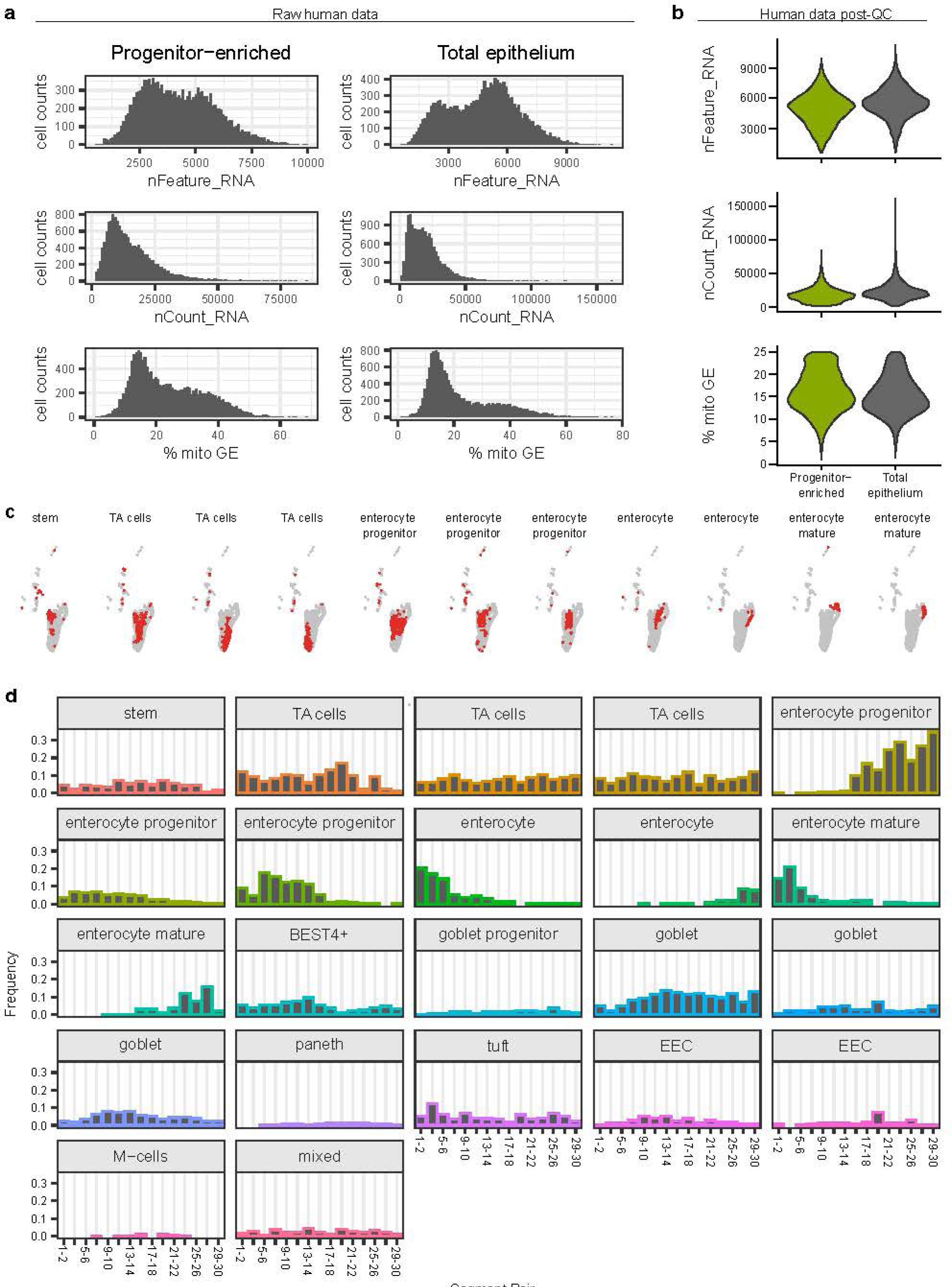
Quality control and initial processing of human scRNAseq data from human subject 2. **a,b** Quality control metrics of data, including number of genes detected (‘nFeature_RNA’), number of unique molecular identifiers detected (‘nCount_RNA’), and percent mitochondrial reads (‘% mito GE’) before (**a**) and after (**b**) processing data. **c** Uniform Manifold Approximation and Projection for Dimension Reduction (UMAP) of total human cells sequenced post-QC, highlighting cell type annotation. **d** Frequency of cells of all epithelial subtypes by segment pair. QC, quality control, mito, mitochondrial; GE, gene expression; ISC, intestinal stem cell; TA, transit amplifying.

**Extended Data Fig. 6.**
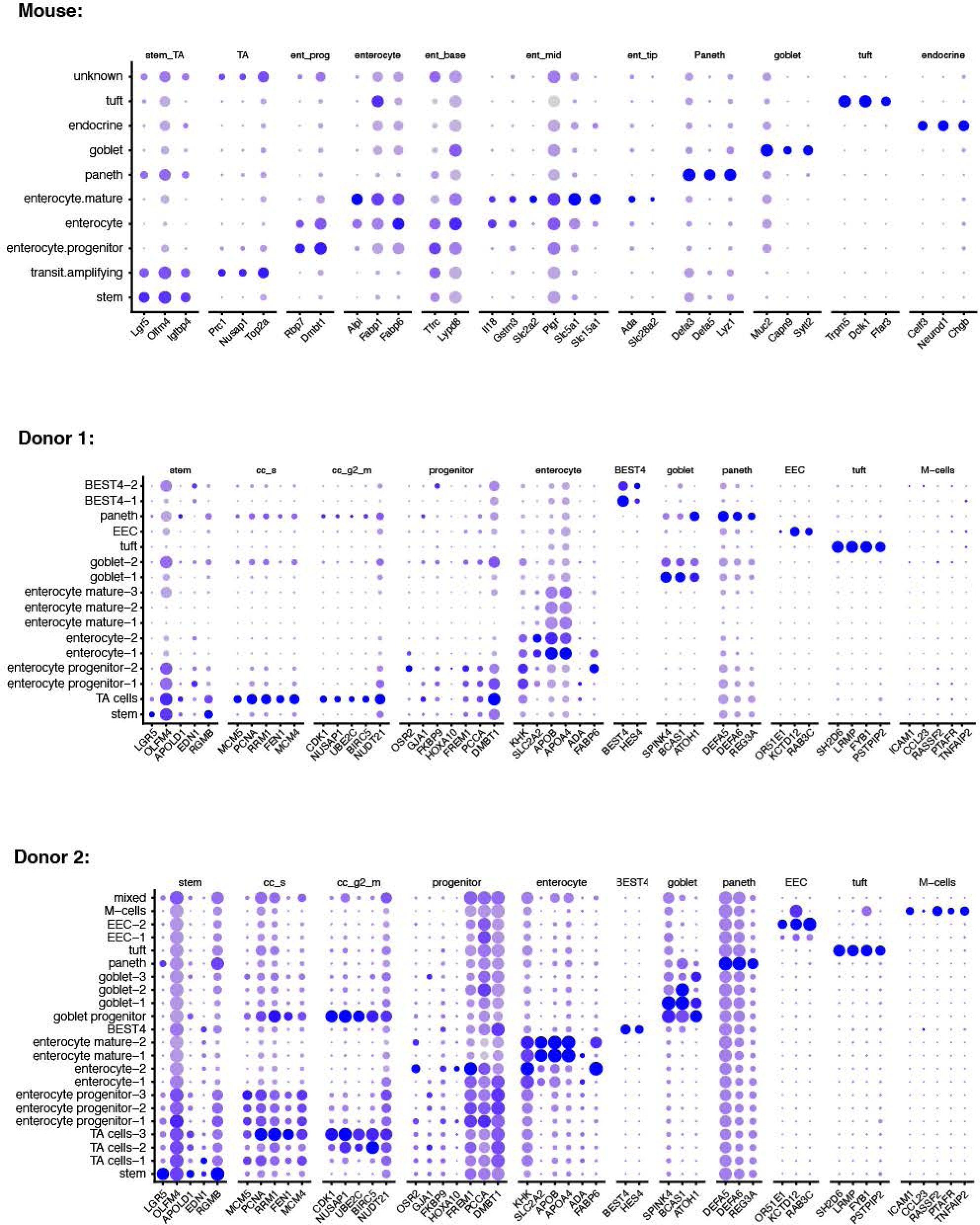
Mouse and human cell type marker genes. **a-c** Dotplots showing expression of cell type marker genes for each cell type sequenced from mouse (**a**) and human donors (**b and c**). See Methods for detailed description of cell type annotation procedures.

**Extended Data Fig. 7.**
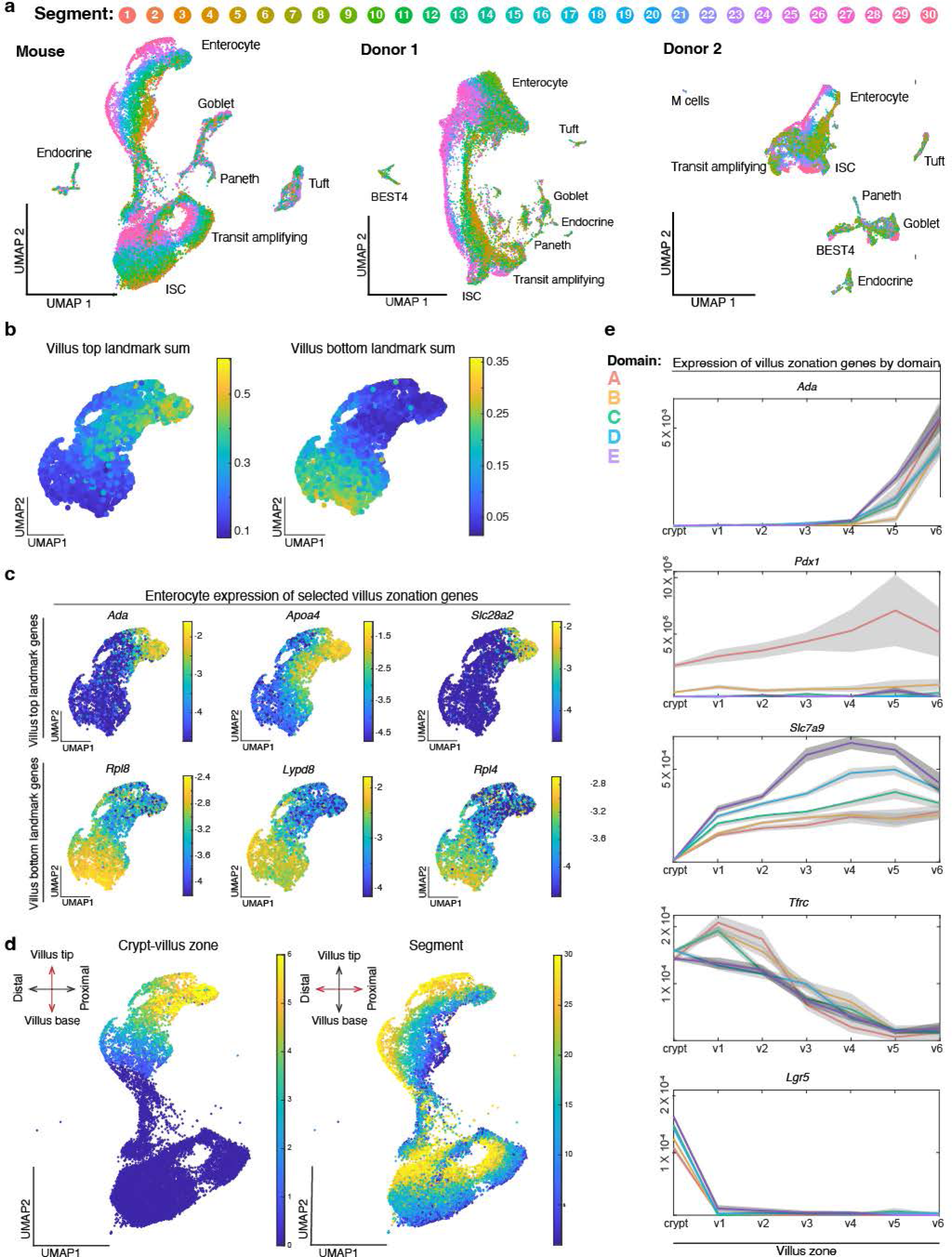
Zonation across multiple axes of the small intestine. **a** UMAP of absorptive lineage cells colored by segment number along the proximal to distal axis in mouse and human donors. Major epithelial cell types are labeled. **b-e** Villus zonation across murine enterocytes. **b** UMAP plots colored according to summed expression of previously reported 14 landmarks of the villus tip (left) or base of villus (right). An equal number of enterocytes were assigned to each of 6 crypt:villus zones, zones 1 - 6. **c** UMAP plots colored according to the expression of select top and bottom villus markers. **d** UMAP plots colored according to villus zonation scores (left) compared to segment positions (right). Villus zonation scores represent the ratio of the summed expression of bottom and top landmark genes. **e** Expression of select villus zonation markers, colored by domain with surrounding grey standard error bands, across crypt:villus zones. UMAP, Uniform Manifold Approximation and Projection.

**Extended Data Fig. 8.**
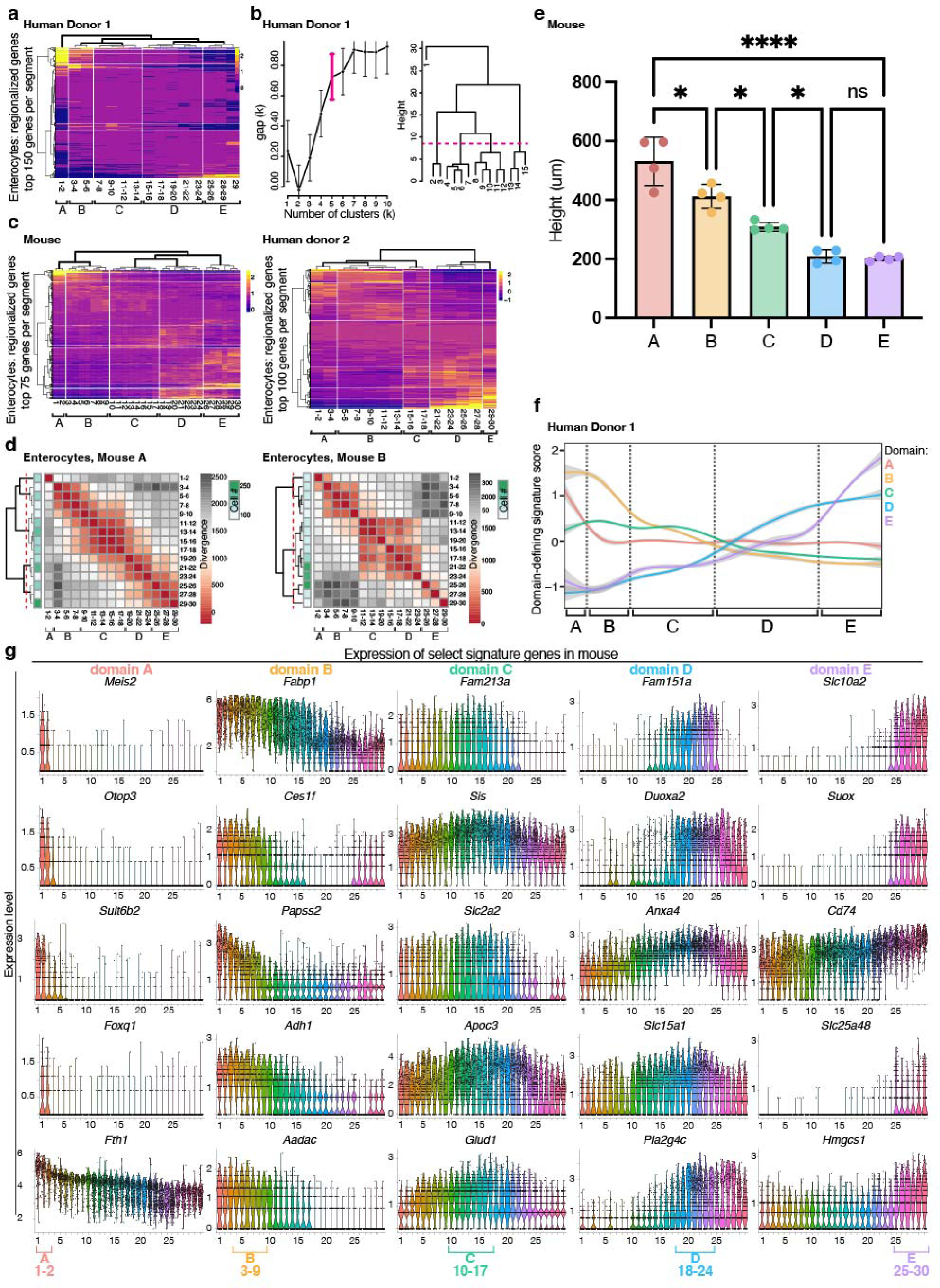
Stability and features of five domains across the mouse and human small intestine. **a** Average expression of the top 150 upregulated genes in enterocytes from human donor 1 in each segment, with segment order and hierarchical clustering based on expression distance between segments. Vertical white lines show the five domains that divide the small intestine, based on: **b** *left:* gap statistics for hierarchical clusters of enterocytes in regional gene expression distance. *Right:* Cuts of dendrogram with optimal cluster number (magenta bracket, left). **c** Most highly regionalized genes expressed by enterocytes in mouse and donor 2 as in Fig. 1f,g but with a smaller number of genes displayed (75-100), as indicated on the y-axis. **d** Jensen-Shannon Divergence between enterocytes from segment pairs across the intestine of each individual mouse, with segment pair order and hierarchical clustering based on divergence values between segments. **e** Average villus height by domain in mouse. Villus base to tip distances were measured for 3-5 villi in each segment, for each of 4 mice. Statistical significance was calculated using one-way ANOVA followed by Tukey’s multiple comparisons test for villus heights across all segments in each domain. *P < 0.05, ****P < 0.0001, ns not significant. **f** Domain-defining gene expression scores for human donor 1, as in Fig. 2c,d, colored by domain with surrounding grey standard error bounds, across intestinal segments. Positions of domain boundaries calculated in **b** are noted with dotted lines and brackets. **g** Expression of key domain marker genes in mouse enterocytes across segments. The segment positions of each domain designation are indicated (bottom).

**Extended Data Fig. 9.**
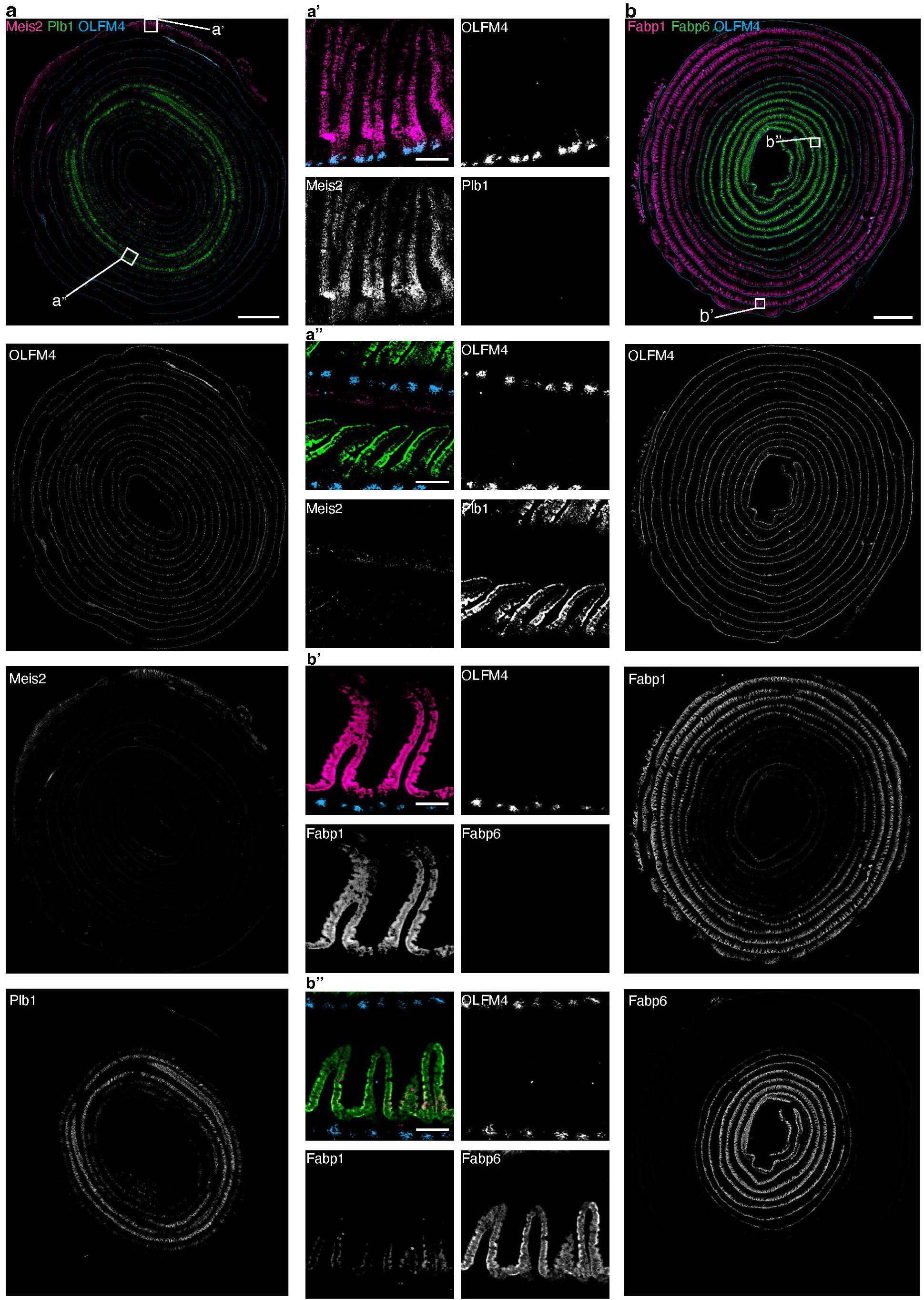
Single-molecule *in situ* hybridization (ISH) validation of key domain markers. **a,b** Full-length murine intestinal tissue coiled from the proximal (outside) end to the distal (inside) end, probed with single-molecule ISH for select marker genes of domains as indicated. Channels are shown both individually and merged with pseudocoloring (as in Fig. 2b,c). White boxes indicate insets. Scale bars are 2 mm, and 100 μm for insets.

**Extended Data Fig. 10.**
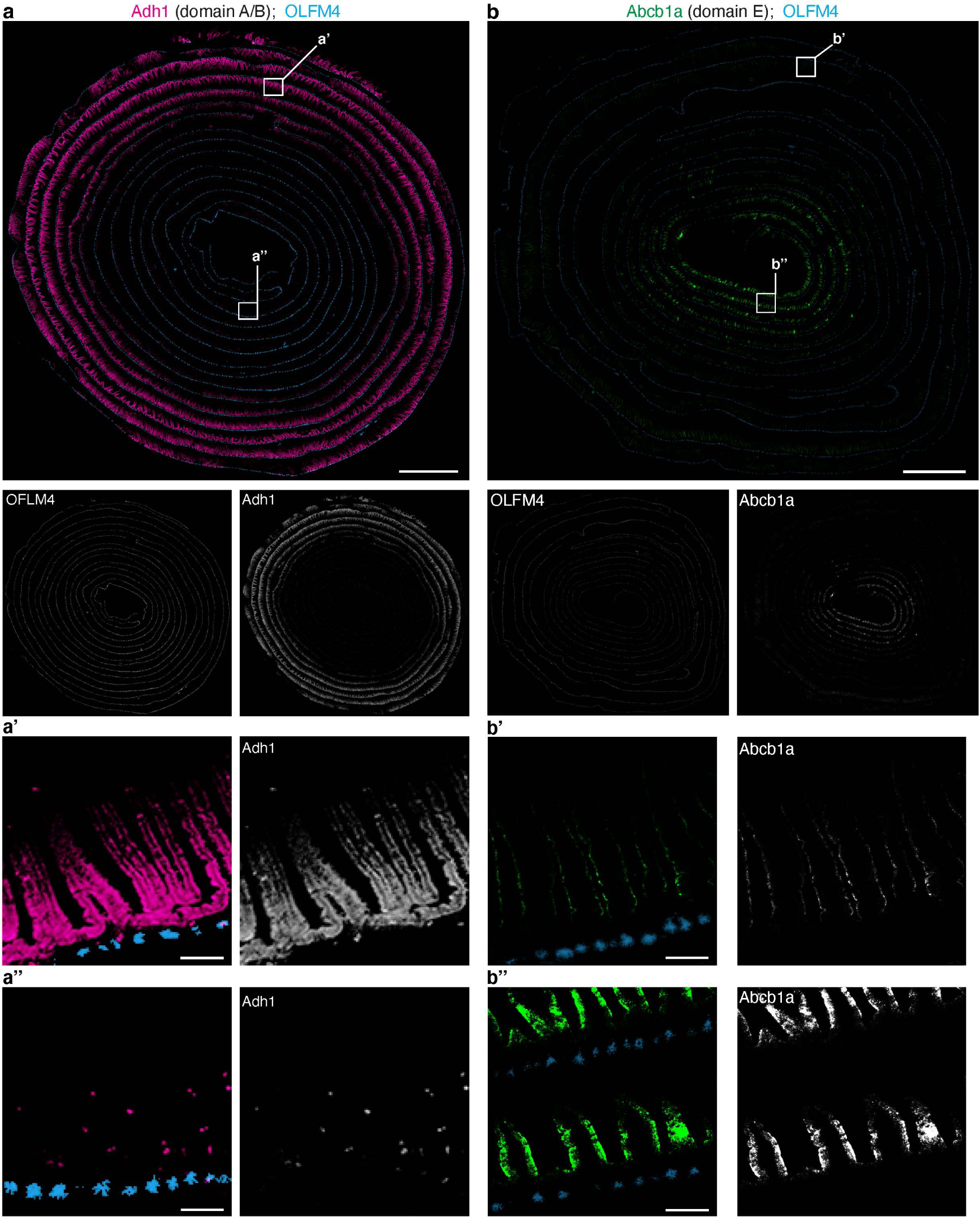
Single-molecule ISH validation of additional domain markers. **a,b** Full-length murine intestinal tissue coiled from the proximal (outside) end to the distal (inside) end, probed with single-molecule ISH for select marker genes of domains as indicated. Channels are shown both individually and merged with pseudocoloring. White boxes indicate insets. Scale bars are 2 mm, and 100 **μ**m for insets.

**Extended Data Fig. 11.**
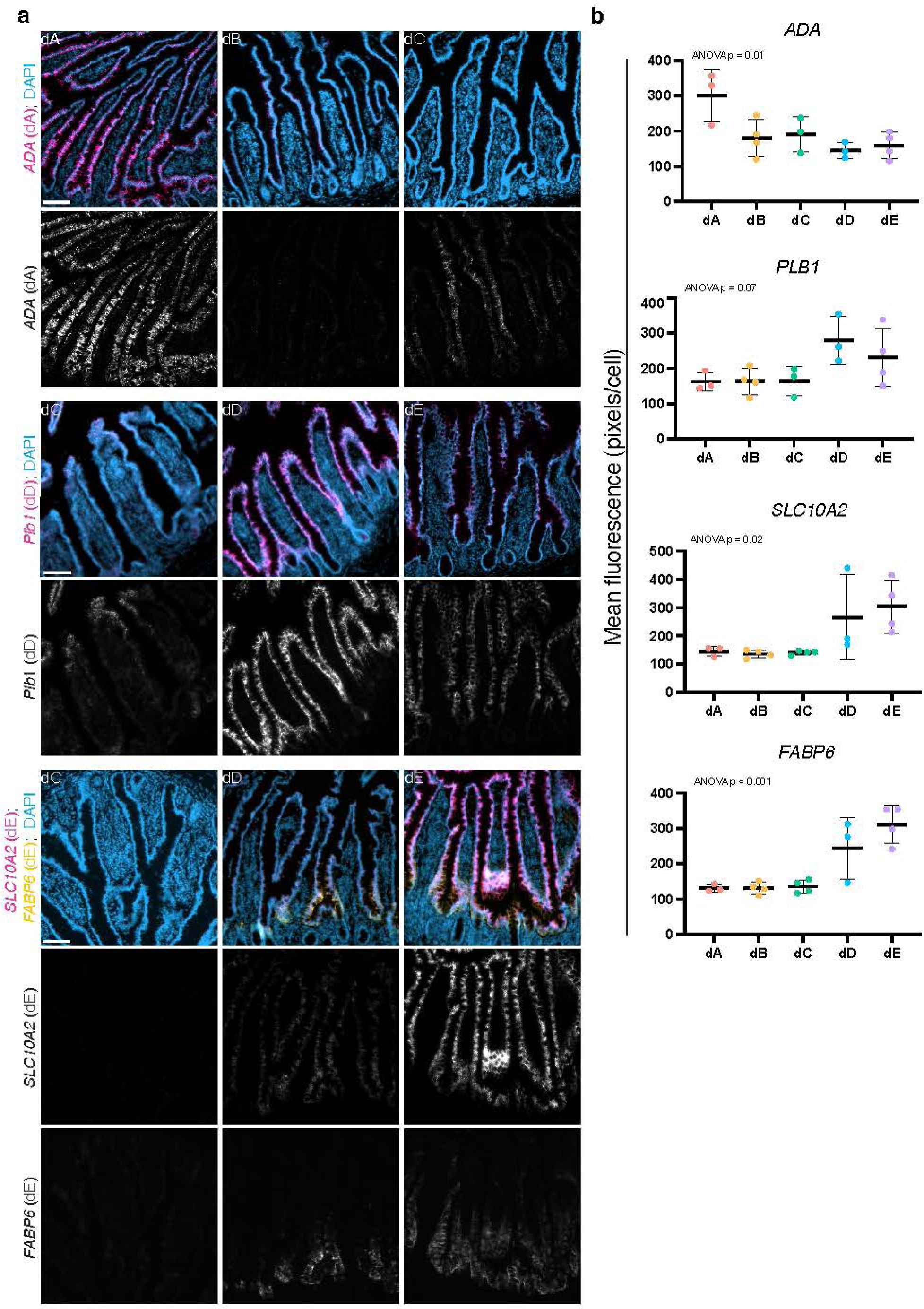
a Single channels of multi-channel images in **Fig. 3b**. Data are human tissue sections from indicated domains probed using single-molecule ISH with domain marker genes. **b** Quantification of mean fluorescence per domain for each donor. n = 3 or 4 donors per domain as indicated by number of datapoints. Scale bars are 100 μm. one-way ANOVA was performed to compare mean fluorescence in each donor by domain, p values for each marker are labeled.

**Extended Data Fig. 12.**
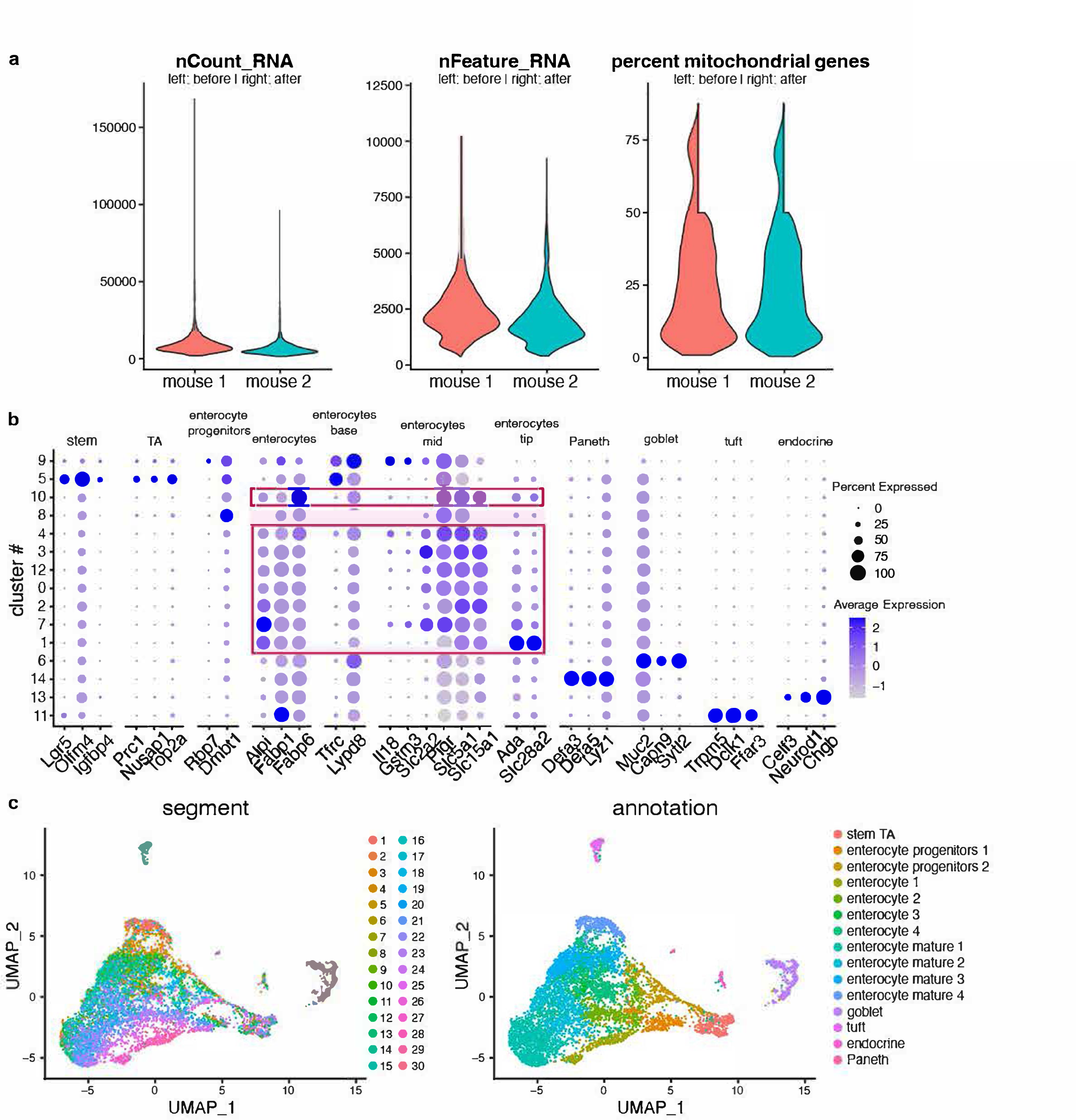
Quality control and initial processing of mouse scRNAseq data used in domain predictions, **Fig. 3d**. **a** Quality control metrics of data including number of unique molecular identifiers detected (‘nCount_RNA’), number of genes detected (‘nFeature_RNA’), and percent mitochondrial genes before and after processing data in each of two mice. **b** Dotplots showing expression of marker genes for each cell type sequenced. Red boxes denote enterocytes, which were the only cell type from these data used for downstream analysis. **c** Uniform Manifold Approximation and Projection for Dimension Reduction (UMAP) of sequenced intestinal epithelial cells post-QC, colored according to segment position (left) and cell type annotation (right). Stem, intestinal stem cell, TA transit amplifying.

**Extended Data Fig. 13.**
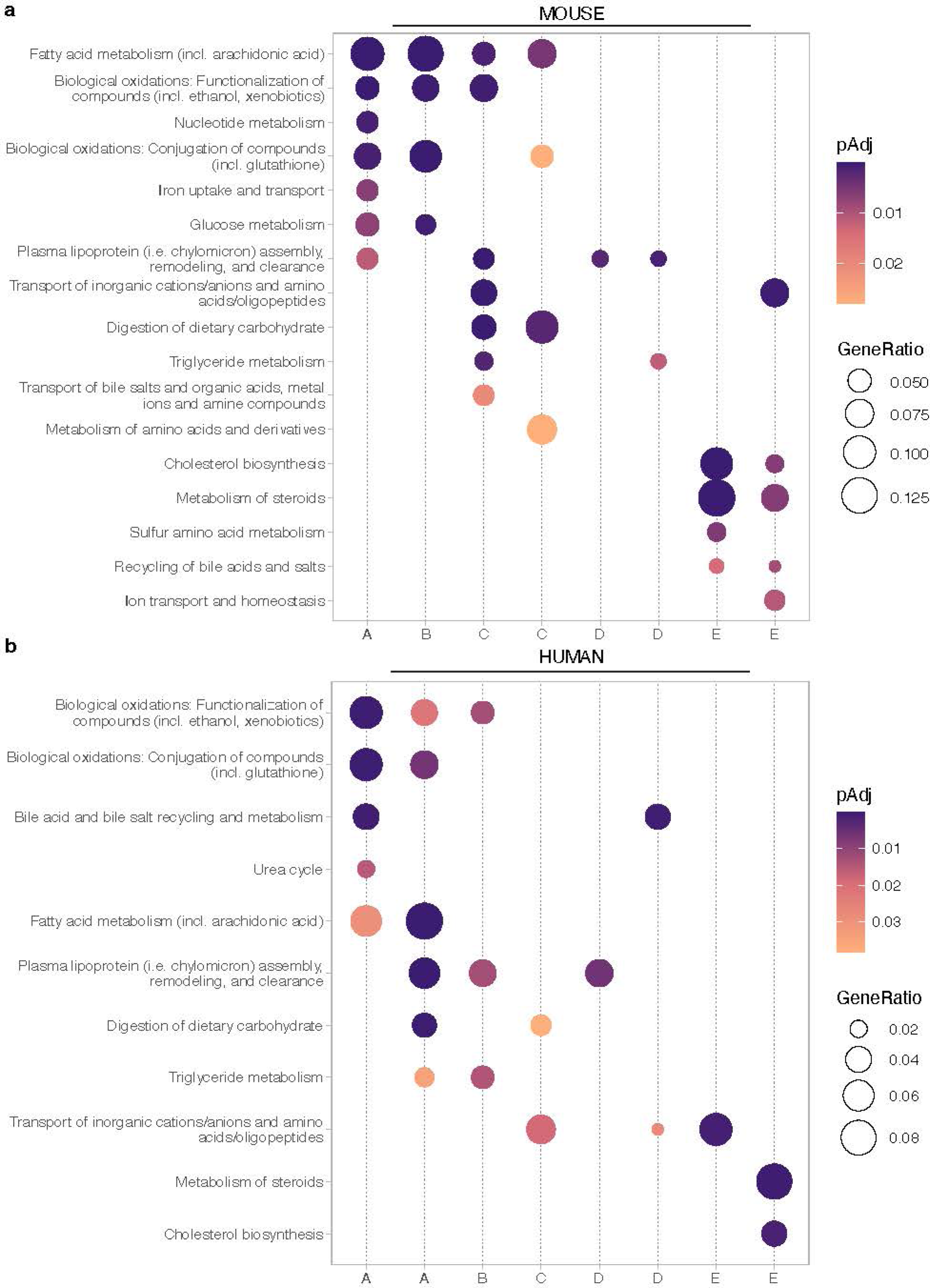
Functional pathways enriched in domain-associated NMF gene modules in mouse and human. **a,b** Selected enriched functional pathways in each NMF gene module displayed in Fig. 2e,f in (**a**) mouse and (**b**) human. All gene modules with a regionally variable expression profile across segments that contained genes that encode aspects of nutrient metabolism are displayed (8 modules per species, dotted vertical lines). Module labels (bottom) are the domain(s) most closely-associated with each module, as determined by regional expression profile and rank of key domain-associated signature genes. Pathways were edited to remove redundancy.

**Extended Data Fig. 14.**
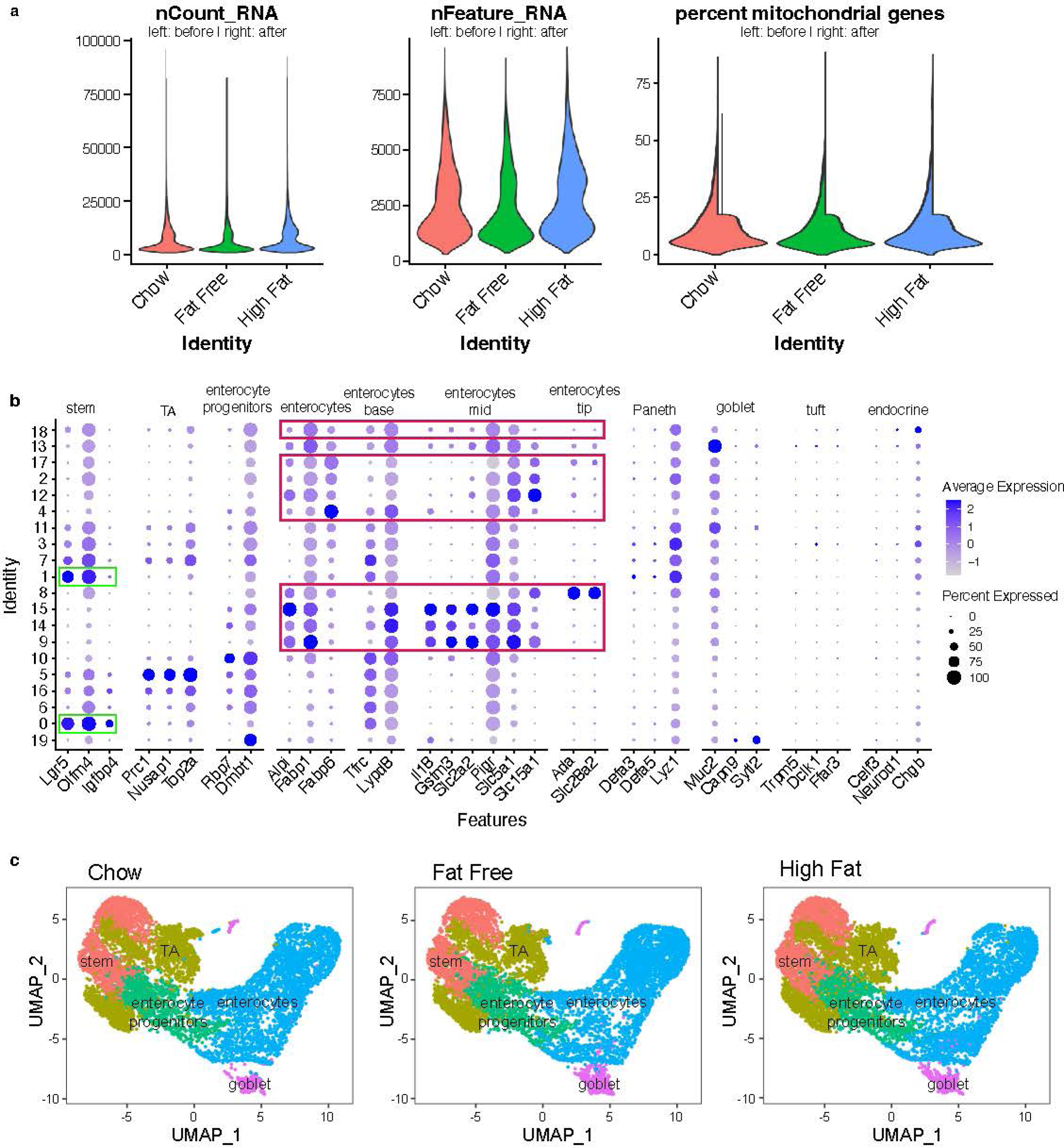
Quality control and initial processing of mouse scRNAseq data used in dietary intervention experiments, **Fig. 4b,c**. **a** Quality control metrics of data including number of unique molecular identifiers detected (‘nCount_RNA’), number of genes detected (‘nFeature_RNA’), and percent mitochondrial genes before and after processing data in each diet group. **b** Dotplots showing expression of marker genes for each cell type sequenced. Red and green boxes denote the cell types analyzed in the study. **c** Uniform Manifold Approximation and Projection for Dimension Reduction (UMAP) of sequenced intestinal epithelial cells post-QC, colored according to segment cell type annotation. Stem, intestinal stem cell, TA transit amplifying.

**Extended Data Fig. 15.**
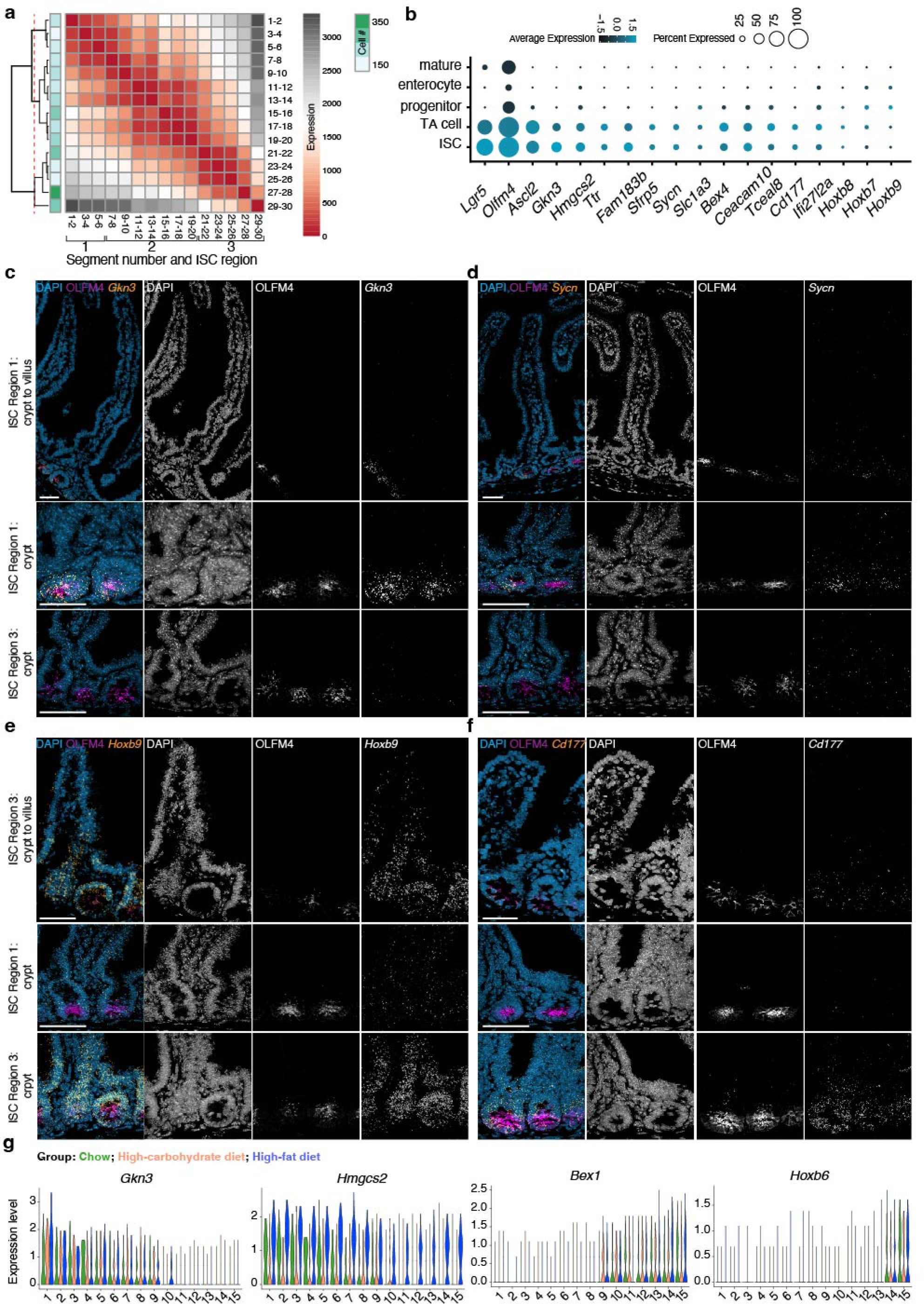
Divisions between regional intestinal stem cells (ISCs). **a** Jensen-Shannon Divergence between ISCs from segment pairs across the intestine, with segment pair order and hierarchical clustering based on divergence values between segments. Dotted red line indicates level of hierarchical tree of domain divisions. **b** Expression of regional ISC marker genes in absorptive lineage cells. Dot color reflects average expression, dot size reflects the percent of cells of each type expressing the marker. **c**–**f** Single-molecule ISH validation of key regional ISC markers. Tile scans displaying full crypt to villus units (top), and crypts from ISC regions 1 and 3 as indicated. Tissue was probed for select regional ISC marker genes as indicated (as in Fig. 5d). Channels are shown both individually and merged with pseudocoloring. Scale bars are 50 μm. **g** Expression of ISC region 1 genes (*Gkn3* and *Hmgcs2*) and ISC region 3 genes (*Bex1* and *Hoxb6*) across ISCs from 15 segments collected from the small intestines of mice fed chow, high-carbohydrate, or high-fat diets as indicated by color. (n = 3 mice per diet group). ‘Mature’ and ‘progenitor’ refer to enterocyte state. ISC, intestinal stem cell; TA, transit amplifying.

**Extended Data Fig. 16.**
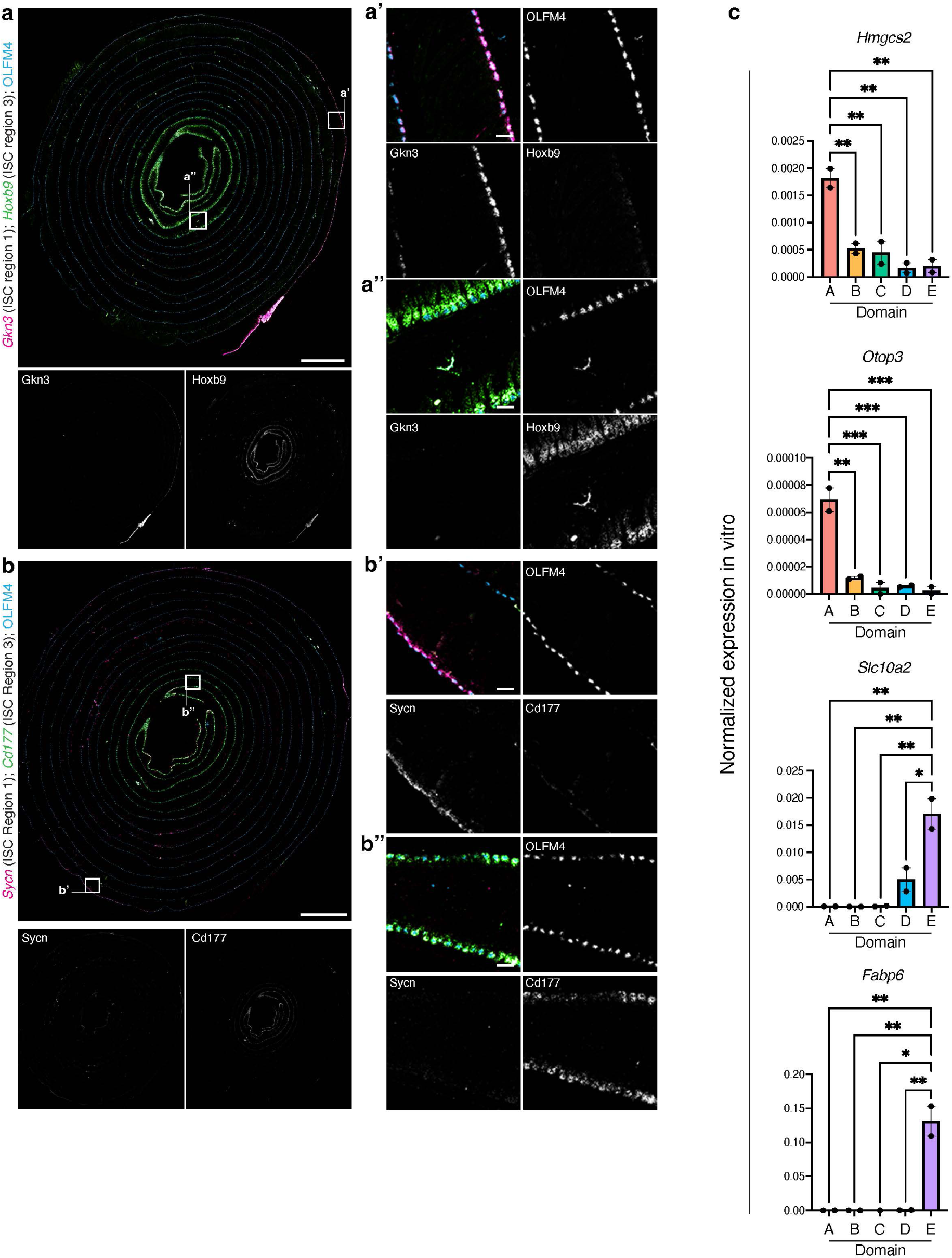
Single-molecule ISH validation of key regional ISC markers. **a,b** Full-length murine intestinal tissue coiled from the proximal (outside) end to the distal (inside) end, probed with single-molecule ISH for select regional ISC marker genes (as in Fig. 5d) as indicated. Channels are shown both individually and merged with pseudocoloring. White boxes indicate insets. Scale bars are 2 mm, and 100 μm for insets. **c** qPCR confirmation of in vitro enrichment of selected Domain A (Hmgcs2, top3) and Domain E (Slc10a2, Fabp6) signature genes in domain A and E-derived organoids respectively, relative to other domain-derived organoid cultures. Regional organoids were cultured for > 1 month and analyzed 5–6 days after passaging. n = 2 organoid lines (biological replicates) per domain.

**Extended Data Fig. 17.**
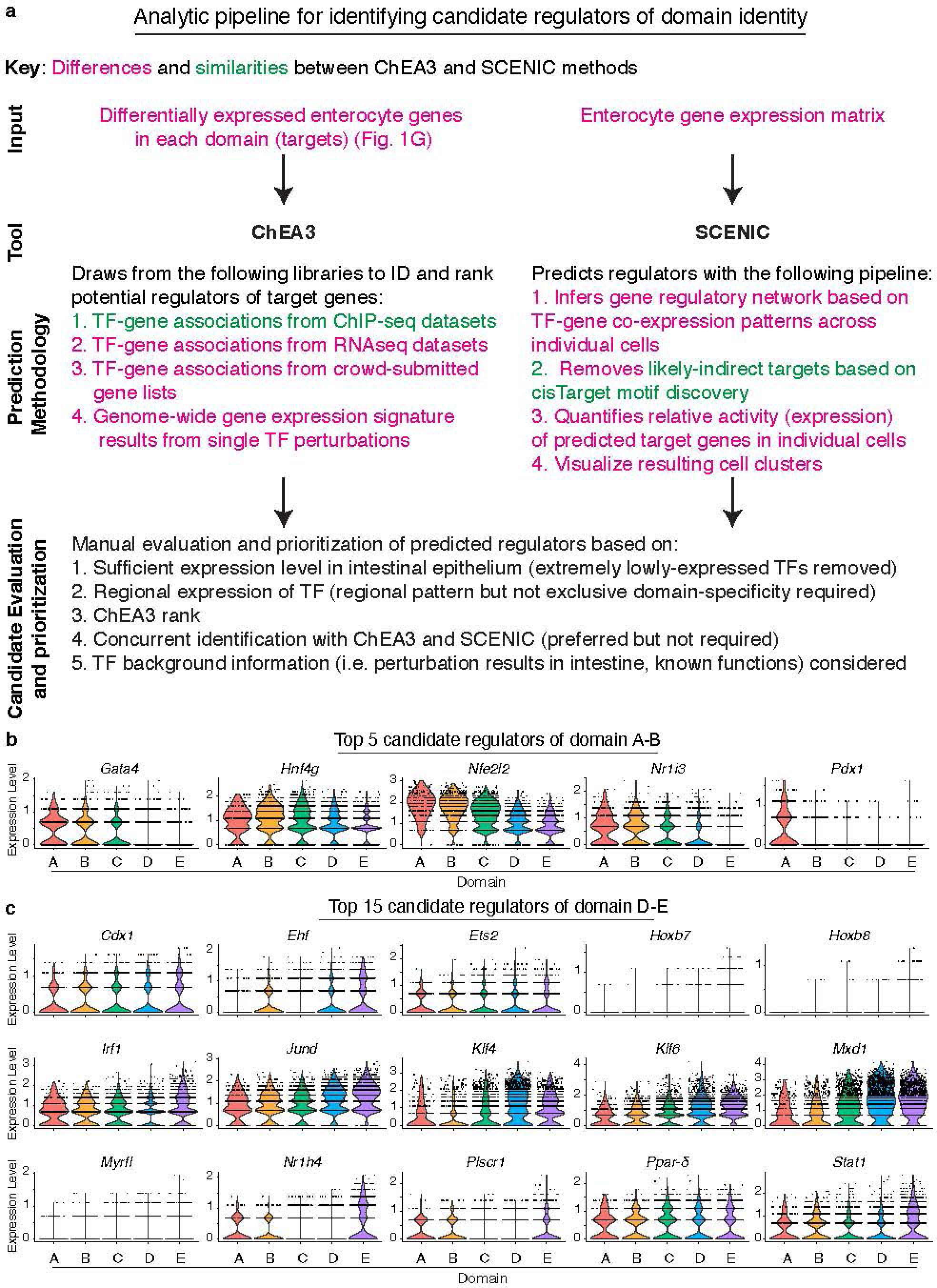
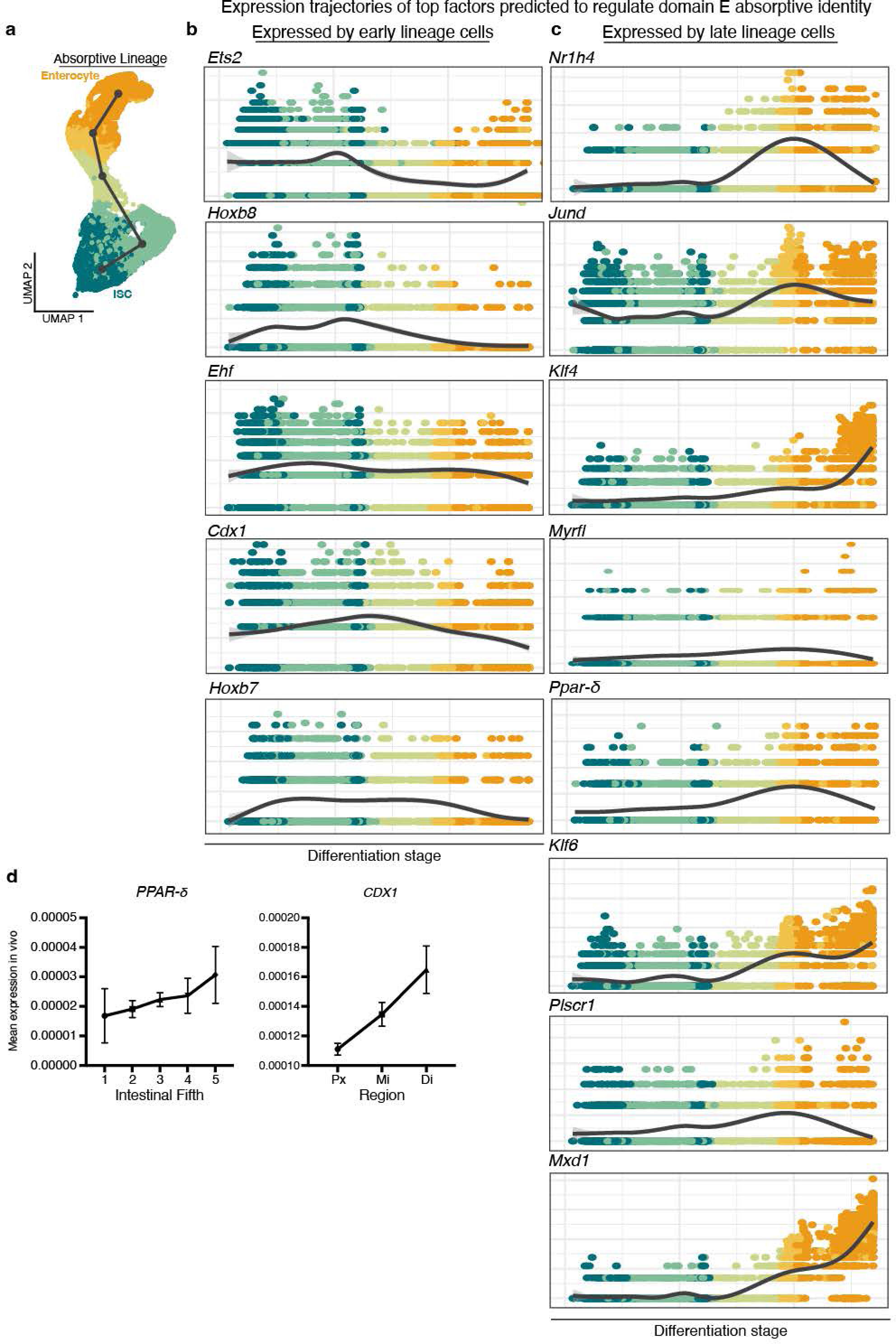
Identification of top candidate regulators of domain identity. **a** Analytic pipeline for predicting regulators of domain identity using gene regulatory inference tools ChEA3 and SCENIC. Methodological distinctions and commonalities between these pipelines indicated in magenta and green, respectively. Criteria for ranking ChEA3 and SCENIC results are described. **b,c** Domain-wise expression levels of 5 candidate regulators of domain A and B identities (**b**) and 15 candidate regulators of domain D and E identities (**c**), identified using the pipeline outlined in **a**.

**Extended Data Fig. 19.**
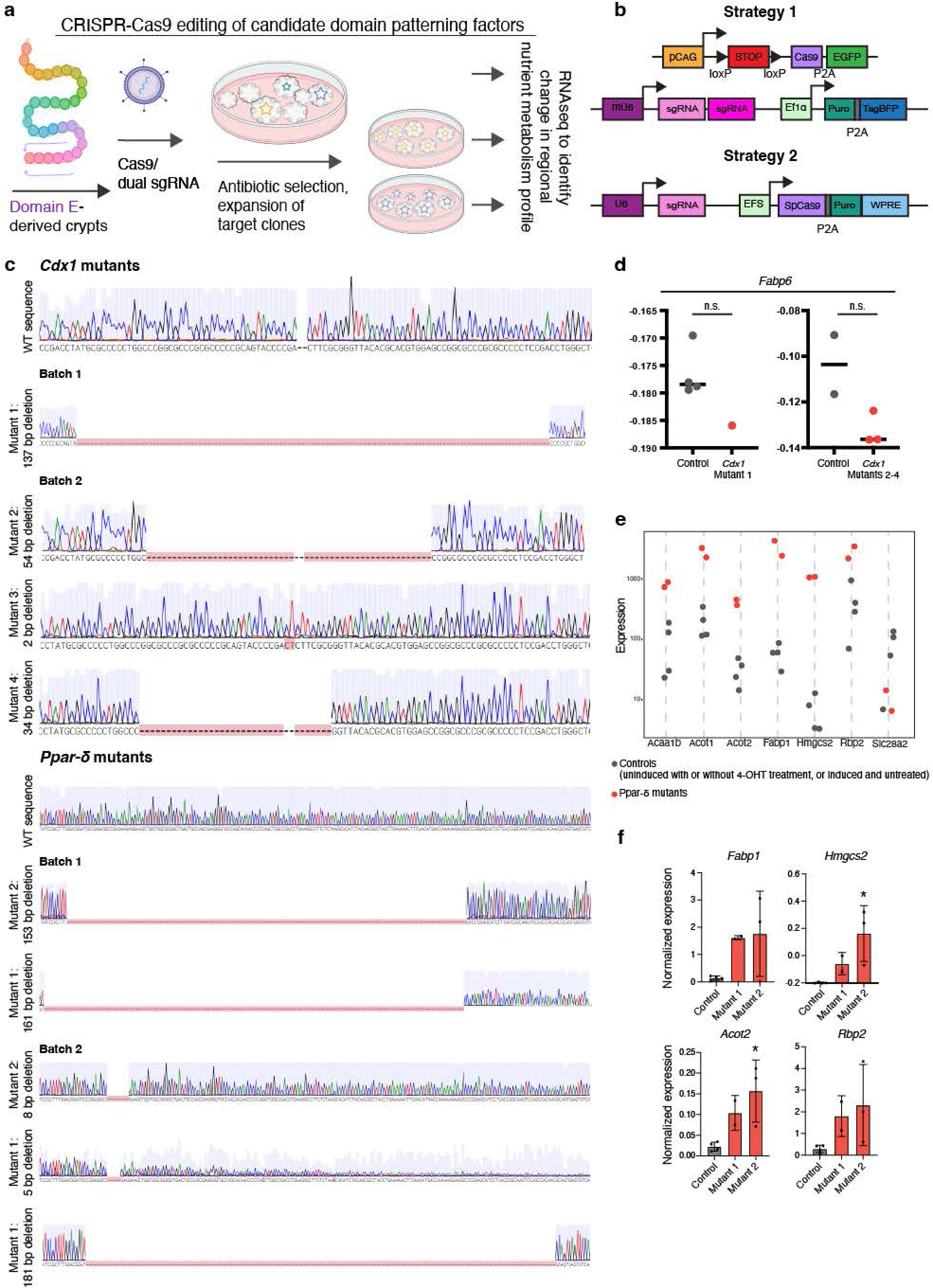
Generation and analysis of *Ppar-ο and Cdx1* mutant domain E organoids. **a,b** Schematics of CRISPR/Cas9 gene targeting strategy. Cas9 endonuclease was encoded in an endogenous genomic locus and 4-hydroxytamoxifen-induced (strategy 1) or delivered by lentiviral vector (strategy 2). Target-specific sgRNAs were delivered by lentiviral vectors (strategy 1 and 2) to induce mutations in the protein coding regions of the target genes. Following mutagenesis, selected clones were expanded and genotyped. Clones containing exclusively deleterious alleles were used for downstream analysis. **c** *Cdx1* mutant organoid sequences from CRISPR editing strategy 1 (**‘**batch 1’, n = 1 mutant line from mouse 1) and 2 (**‘**batch 2**’,** n = 3 unique mutant lines from mouse 2), and *Ppar-ο* mutant organoid sequences from editing strategy 1 (**‘**batch 1’, n = 2 mutant line from mouse 1) and 2 (**‘**batch 2**’,** n = 3 unique mutant lines from mouse 2). Indel mutations are specified. **d** Trend towards decreased expression of *Fabp6* in Cdx1 mutant lines in both batches of mRNAseq expression data from editing strategies 1 and 2, which could not be merged. Line represents median. **e** Expression of differentially expressed genes in individual *Ppar-ο* mutant organoid lines from batch 1 mutants (red dots) and control organoid lines (black dots). Batch 2 expression data of these and other DEGs in Fig. 6e,g. **c** Normalized mRNA levels of select DEGs of interest in *Ppar-ο* mutant organoids, validated with real time PCR. *P < 0.05 calculated using one-way ANOVA with Tukey’s multiple comparison test. Data are mean ± SD (2-3 technical replicates per line). bp, base pair; DEGs, differentially expressed genes, n.s. not significant.

