## Supplemental Table 9 for "Epithelial zonation along the mouse and human small intestine defines five discrete metabolic domains"

Supplementary Table 7: Primer Sequences

qPCR primers

| **Gene** | **Primer Sequence 5’-3’** |
| --- | --- |
| *Ppar-δ* | **For** CTGCTCACTGACAGATGAAGAC  **Rev** GTGGCTGTTCCATGACTGA |
| *Cdx1* | **For** GACGCCCT ACGAATGGATG  **Rev** GTACCGGCTGTAGTGAAACTC |
| *Fabp1* | **For** TTGCCACCATGAACTTCTCC  **Rev** CCCTTGATGTCCTTCCCmC |
| *Hmgcs2* | **For** CAGCAGTGACAAACAGAACAAC  **Rev** GAGCCTTGTCT ACATCCTTGT |
| *Acot2* | **For** GGAACACCATCTCCTACAAGG  **Rev** AGCTTCCACGACATCCAAG |
| *Rbp2* | **For** CTTTGAAGGCTACATGAAGGC  **Rev** GTGCTGTTGGTTTTCGTCTTG |
| *Otop3* | **For** TGACCAATGACTCCATGCAC  **Rev** AGGAAAAGACAGCATCTCAGG |
| *Slc10a2* | **For** TGTTCACCTTCCCACTCATC  **Rev** GTCCATTTCATTGTCTGTTTTCTCT |
| *Fabp6* | **For** CCCAACTATCACCAGACTTCG  **Rev** GCCAGCCTCTTGCTTACG |

sgRNAs

| **Target** | **Cassette Vector** | **sgRNA sequence** |
| --- | --- | --- |
| *Ppar-δ* | pU6sgRNA-EF1alpha-puro-T2A-BFP | 5’ **For**  TTGGCCTCCGGCATCCGTCCAAAGGTTTAAGAGC  3’ **Rev** TTAGCTCTTAAACCTTTGGACGGATGCCGGAGGCCAACAAG |
| *Ppar-δ* | pMJ117 | 5’ **For**  ATGGACATGACCAAAAAGAAGGCCGTTTCAGAGC  3’ **Rev** TTAGCTCTGAAACGGCCTTCTTTTTGGTCATGTCCATGTTT |
| *Cdx1* | pU6sgRNA-EF1alpha-puro-T2A-BFP | 5’ **For**  TTGGGCGTGTAACCCGCGAAGTCGGTTTAAGAGC  3’ **Rev** TTAGCTCTTAAACCGACTTCGCGGGTTACACGCCCAACAAG |
| *Cdx1* | pMJ117 | 5’ **For**  ATGGCGAATGGATGCGGCGCAGCGGTTTCAGAGC  3’ **Rev** TTAGCTCTGAAACCGCTGCGCCGCATCCATTCGCCATGTTT |

Cassette Primers

| **Cassette Vector** | **Primer Sequence 5’-3’** |
| --- | --- |
| pU6sgRNA-EF1alpha-puro-T2A-BFP | **For** tggttagtaccgggccc  **Rev** GCTGAGTGTAGATTCGAGCaaaaaaagcaccgactcg |
| pMJ117 | **For** GCTCGAATCTACACTCAGC  **Rev** cgcctaatggatcctagtactcg |

Target gene PCR and sequencing primers

| **Gene** | **Application** | **Primer Sequence 5’-3’** |
| --- | --- | --- |
| *Ppar-δ* | PCR | **For** AGTCAGCAAAGGCTCTCATCAT  **Rev** CTTCTCTGCCTGCCACAGT |
| *Ppar-δ* | Sequencing | **For** TCTCATCATGGAGCTGCTTCC  **Rev** GAAAACCAGGCCCATCAGA |
| *Cdx1* | PCR | **For** CAA GGA CTC CCC CGT GTA C  **Rev** CTCTCGAGAAGCCAATCAGGG |
| *Cdx1* | Sequencing | **For** TGGGCCCTCCGACCTATGC  **Rev** AGAGAAGGGTCCTTACCGC |
